# Identification of candidate ciliary microtubule-associated proteins using CEP41 proximity labeling

**DOI:** 10.64898/2026.10.27.762278

**Authors:** Xiaobo Sun, Yan Liang, Jinghua Hu

**Author notes:** Correspondence (J.H.). 200 First Street SW, Rochester, MN 55905. These authors contributed equally (Xiaobo Sun, Yan Liang). Senior author (Jinghua Hu).

## Abstract

Primary cilia are microtubule-based signaling organelles whose dysfunction results in a wide spectrum of ciliopathies. Extensive studies have established the functional and clinical importance of ciliary microtubule associated proteins (ciliary MAPs), but the repertoire of specialized ciliary MAPs remains incompletely defined. Here, we used the ciliopathy-associated MAP CEP41 as a BioID bait to probe proteins proximal to ciliary microtubules. Quantitative proteomics and cilia-focused prioritization identified a ciliary MAP-enriched protein set, revealed a poorly characterized Septin-9 isoform, and supported C7orf57/FCAP33, CCDC34, and PSRC1 as novel ciliary MAPs. CCDC34 and PSRC1 did not affect cilia architecture or ciliation, indicating roles in structure-independent ciliary functions, whereas C7orf57 overexpression increased cilium length. Microtubule association analysis reveals C7orf57 as a paclitaxel sensitive MAP that associates with microtubule through its N-terminal regions. Together, these findings expand the ciliary MAP repertoire and provide a resource for prioritizing candidates for future studies of cilia biology and ciliopathies.

## INTRODUCTION

The primary cilium is a sensory organelle that senses environmental cues and transduces diverse signals through pathways including Sonic Hedgehog (SHH) and various G protein-coupled receptor signalings.^1^ Dysfunction of primary cilia results in a broad spectrum of congenital syndromic diseases, collectively termed ciliopathies.^2^ Structurally, the primary cilium contains a membrane-enclosed axoneme, which is an array of nine doublet microtubules stemming from the plasma membrane-docked mother centriole (the basal body).^3^ As a microtubule-based organelle, the primary cilium recruits a distinct pool of ciliary microtubule-associated proteins (MAPs) that maintain axonemal architecture, ciliary homeostasis, and signaling capacity. The motors kinesin-2 and dynein-2 associate with Intraflagellar Transport (IFT) complexes to mediate cargo transport along the axoneme.^4^ Ciliary MAPs have reported functions in regulating ciliary architecture, ciliogenesis, and cilium length.^5–7^ The non-motile motor KIF7, for example, organizes the ciliary tip compartment and subsequent cilia-mediated SHH signaling.^8^ The physiological importance of ciliary MAPs is underscored by variants that cause ciliopathies, including Joubert syndrome, Meckel-Gruber syndrome, and Retinitis pigmentosa.^9–13^ Despite these advances, the full composition of the ciliary MAP repertoire and the mechanisms by which individual MAPs shape cilium architecture and function remains incompletely understood.

Tubulin posttranslational modifications (PTMs) contribute to “tubulin code” to diversify microtubule structures so they can be recognized by cellular effectors to alter microtubule stability or function.^14,15^ Among these PTMs, polyglutamylation occurs predominantly along the ciliary axoneme and neuronal axons, but not in the cytosol.^16,17^ Functionally, axonemal glutamylation has been shown to directly regulate ciliary membrane protein positioning and SHH signaling.^18,19^ Moreover, dysfunctions of several ciliopathy-associated proteins result in decreased ciliary polyglutamylation level.^8,18,20,21^ These observations motivated a search for additional regulators of this tubulin PTM.

To identify candidate ciliary MAPs and potential regulators of axonemal glutamylation, we selected a well-characterized ciliary MAP as a bait to probe its proximal proteome. Ideally, this bait would be broadly distributed across ciliary compartments to maximize chances of finding new ciliary MAPs at both the basal body and axoneme. It is also preferable that this bait is an axoneme polyglutamylation regulator so that other regulators may be detected in its proximal proteome. Joubert syndrome protein CEP41 is a MAP that localizes to both basal bodies and the axoneme.^21,22^ CEP41 regulates axonemal glutamylation with comparatively limited effects on ciliary architecture.^21^ We therefore selected CEP41 as the bait for our proximity proteomics study.

In this study, we used CEP41 based proximity proteomics together with cilia-focused prioritization to generate a curated resource of candidate ciliary MAPs. The resulting dataset was enriched for established ciliary MAPs, microtubule inner proteins, and regulators of tubulin PTMs. Co-localization analyses supported CCDC34, PSRC1, and C7orf57 as novel ciliary MAPs. In addition, we discovered a poorly characterized isoform of Septin-9, which is an established ciliary MAP with multiple well-characterized functional isoforms.^23,24^ Although preliminary RNAi screen on selected candidates did not yield high-confidence axoneme polyglutamylation regulators yet, functional screens on cilia structure showed that C7orf57 overexpression results in abnormally long primary cilia with increased microtubule resistance to nocodazole. Truncation analysis determined that C7orf57 is a paclitaxel sensitive MAP which associates with microtubules through its N-terminal regions. This study can serve as a resource for expanding the ciliary MAP landscape and prioritizing future ciliary MAPs and candidate ciliopathy genes identification.

## RESULTS

### CEP41 is selected as a bait for proximity labeling of candidate ciliary MAPs

The bait for proximity labeling of ciliary MAPs should ideally be broadly distributed along the basal body-axoneme axis so that candidate MAPs across ciliary compartments can be detected. Consistent with previous findings, endogenous CEP41 showed broad localization in primary cilia of both hTERT-RPE1 and RCTE cells (Figure 1A and 1B).^21^ Remarkably, despite being a MAP, endogenous CEP41 shows no detectable binding with cytoplasmic microtubules (Figure 1A and 1B). Exogenously expressed human CEP41-mCherry protein decorated cytoplasmic microtubules only at high expression levels but was predominantly ciliary at low expression levels (Figure 1C). This preferential binding of CEP41 to ciliary microtubules allows us to generate a low-expressing CEP41-BioID cell clone to limit background noise from non-specific binding of CEP41 to cytoplasmic microtubules. After single clone selection, we detected no significant difference in the proportion of ciliated cells or cilium length between Myc-BirA* (BirA R118G mutant; Clone 1G9) and hCEP41-myc-BirA* (Clone 2F9) RCTE cells (Figure 1D and 1E). Western blot analysis showed successful protein biotinylation in both cell lines, with clone 2F9 showing only slightly higher exogenous CEP41-myc-BirA* fusion protein level as compared to endogenous CEP41 level (Figure 1F). Immunofluorescence showed strong ciliary and minimal cytoplasmic microtubule staining of the CEP41-myc-BirA* fusion protein in clone 2F9. Nevertheless, streptavidin staining remained detectable on cytoplasmic microtubules, possibly reflecting residual cytoplasmic biotinylation by CEP41-myc-BirA*. Despite the undesired background staining on cytoplasmic microtubules in clone 2F9, the Myc-BirA* control showed diffuse biotinylation, whereas CEP41-myc-BirA* produced a clear and strong ciliary signal in clone 2F9 (Figure 1G).

**Figure 1.**
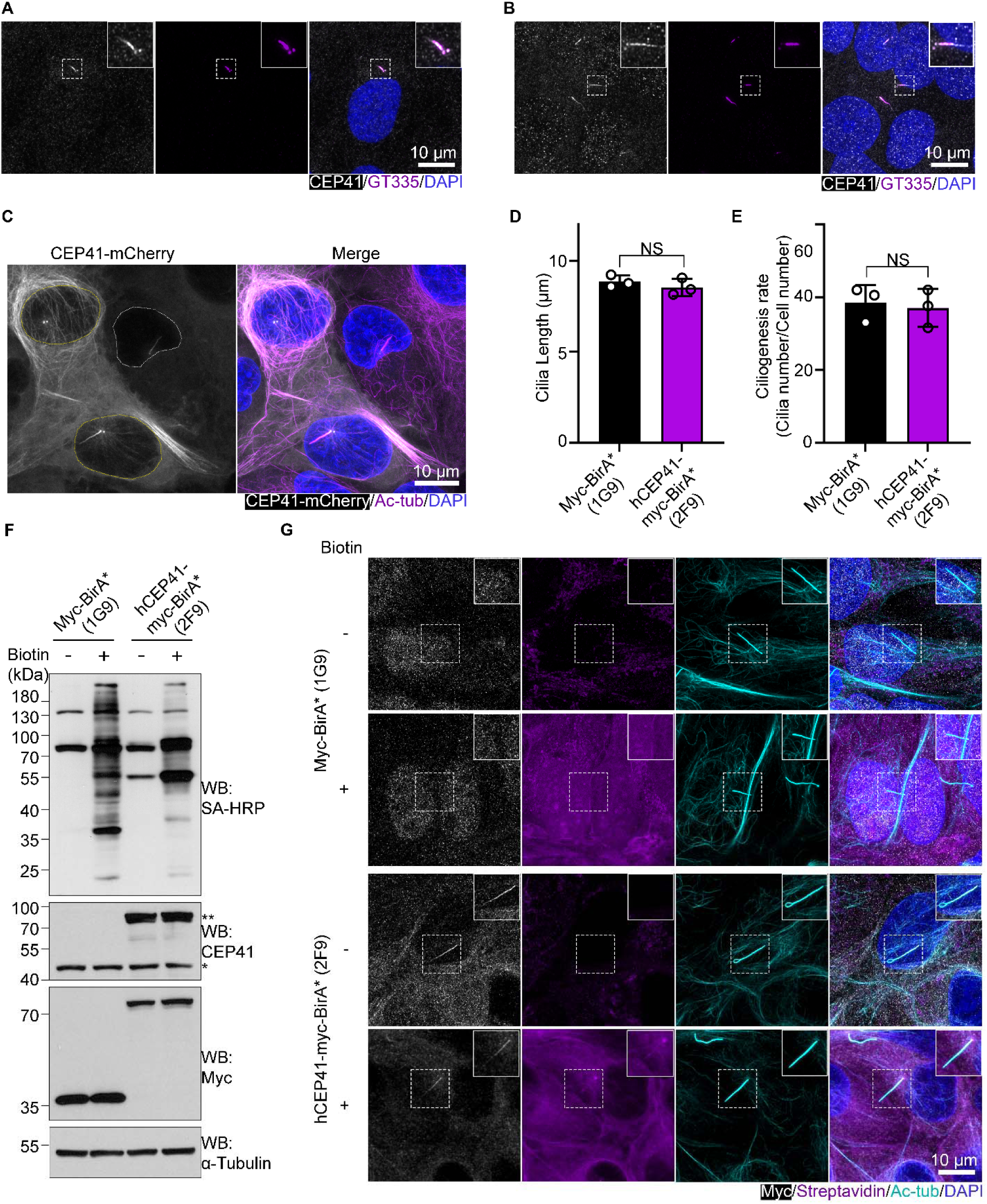
CEP41 is selected as a bait for proximity labeling of candidate ciliary MAPs. Co-localization of CEP41 with cilia marker GT335 in (A) hTERT-RPE1 and (B) RCTE cells. (C) Overexpression of CEP41-mCherry in RCTE cells. Immunofluorescence was performed to co-stain CEP41-mCherry and cilia marker acetylated-tubulin (Ac-tub). Cells with yellow nuclear contour represent high CEP41-mCherry expression. The cell with white nuclear contour represents low CEP41-mCherry expression. RCTE cells stably expressing either Myc-BirA* or hCEP41-myc-BirA* were derived from two single clones 1G9 and 2F9 respectively. Using Ac-tub as cilia marker, (D) cilia length and (E) ciliogenesis rate were compared between two clones. Significance was tested by two-tailed t-test with Welch’s correction. Nonsignificance is abbreviated as NS with p value larger than 0.05. Error bars denote ±SD. Western blot (F) and immunofluorescence (G) validation of protein biotinylation and construct expression in 1G9 and 2F9 clones. hCEP41: Human CEP41. All experiments were performed in triplicate.

### CEP41 proximity proteomics defines a curated set of 40 candidate ciliary MAPs

To define the cilia-enriched CEP41 proximity proteome, we first induced ciliogenesis followed by biotin addition in clones 1G9 and 2F9. After streptavidin pulldown, we confirmed that α-tubulin and GT335 reactive glutamylated tubulin were enriched in 2F9 samples as compared with 1G9 samples (Figure 2A and 2B). Following on-bead digestion and LC-MS/MS, 365 proteins were enriched at least twofold with an adjusted p-value of less than 0.05 in 2F9 relative to 1G9 samples (Data S1). Gene Ontology (GO) analysis confirmed that enrichment of ciliary and microtubule-related proteins in CEP41 proximity proteomes (Figure 2C).

**Figure 2.**
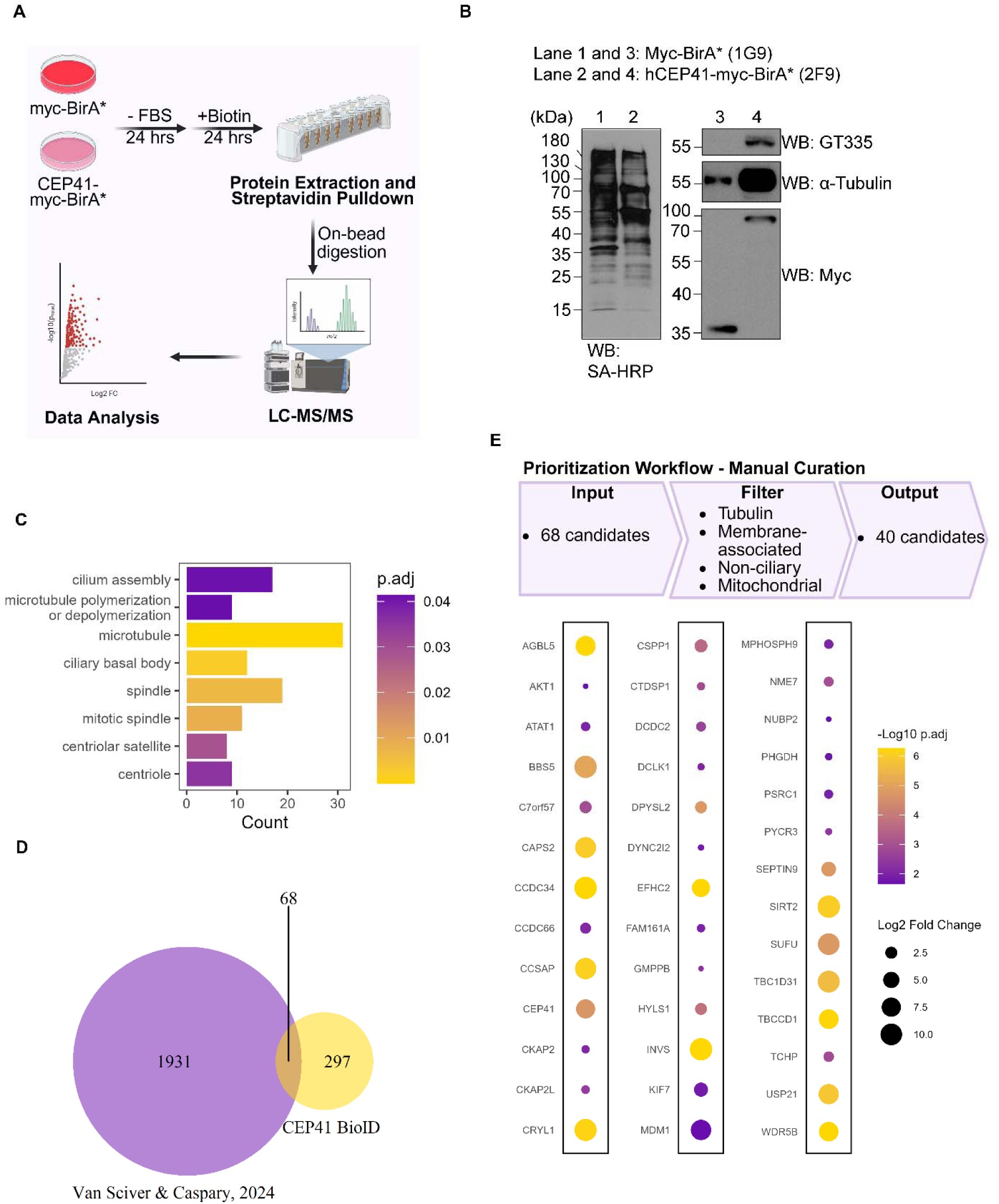
CEP41 proximity proteomics defines a curated set of 40 candidate ciliary MAPs. (A) Schematics showing workflow of characterizing CEP41 proximity proteomes using BioID. (B) Validation of enriched biotinylated proteins in Myc-BirA* (1G9) and CEP41-myc-BirA* (2F9) samples after protein extraction and streptavidin pulldown. (C) Gene Ontology (GO) analysis reveals statistically significant cilia and microtubule related GO terms. The color of the graph represents adjusted p value. The x-axis represents the number of proteins found in each GO term. (D) Venn diagram of cross-reference between unified dataset from van Sciver & Caspary, 2024 and significantly enriched proteins found in CEP41 BioID dataset.^25^ 68 proteins were found to be overlapped in two datasets. (E) The upper panel shows prioritization workflow of manual curation to select candidates for downstream functional analysis. Dot plot shows the list of 40 proteins. The size of the dot represents fold change of enrichment in 2F9 over 1G9 samples. The color represents adjusted p value.

To prioritize candidate ciliary MAPs, we first cross-referenced the CEP41-enriched dataset with a unified resource combining ciliary phenotypic datasets from model organisms, genetic and cilia-related proteomics and transcriptomic datasets.^25^ Because this resource includes both established ciliary proteins and candidates with limited validation, we believe that the overlap could yield a focused but deliberately cilia-biased candidate set with the potential to find novel ciliary MAPs. Cross-referencing the two datasets identified 68 overlapping candidates (Figure 2D). We then excluded tubulins, membrane-associated proteins, experimentally validated non-ciliary proteins, and likely mitochondrial contaminants.^26^ Manual curation yielded a prioritized set of 40 CEP41-proximal proteins for downstream ciliary MAP screening (Data S2 and Figure 2E).

Of note, our prioritized set contained eight established ciliary MAPs (i.e. INVS, MDM1, AGBL5, CEP41, KIF7, CSPP1, DCDC2, and FAM161A) implicated in sensory ciliopathies.^10,12,13,21,27–30^ Eight additional ciliary MAPs (i.e. SIRT2, USP21, Septin-9, DPYSL2, CCDC66, NME7, ATAT1, and CKAP2L) in the CEP41 proximity proteome have been reported to regulate primary cilia homeostasis and functions.^5,24,31–36^ Recent *in situ* cryo-electron tomography identified CEP41 as a microtubule inner protein (MIP) in mouse ependymal basal bodies and cilia.^37^ Consistent with this finding, our curated CEP41 proximal protein set also contained numerous validated or predicted ciliary MIPs, including MDM1, CAPS2, EFHC2, CSPP1, TCHP and NME7, in our curated CEP41 proximal proteomes.^38–40^ We also recovered three ciliary MAPs (i.e. AGBL5, CCSAP, and KIF7) that were previously shown to regulate tubulin glutamylation (Data S2).^8,41,42^ Recovery of established ciliary MAPs, MIPs, and glutamylation regulators supports the biological relevance of the dataset and its potential to uncover novel ciliary MAPs and regulators of axoneme glutamylation.

### Candidate screening supports CCDC34, PSRC1, and C7orf57 as candidate ciliary MAPs

Among the 40 CEP41-proximal proteins, eleven were selected for localization screening because their roles in primary cilia were poorly characterized at the start of the study. We generated stable RCTE cell lines expressing each of the eleven Flag-tagged candidates, assessed ciliary localization, and then evaluated co-localization with the cytoplasmic microtubule arrays (Figure 3A). We confirmed that our Flag antibody did not detect either cilia or microtubules (Figure S1A and S1B). Tagged CRYL1 and GMPPB showed diffuse cytoplasmic and nuclear localization, whereas PHGDH and PYCR3 showed only cytoplasmic signals (Figure S1E and S1H-S1J). In addition, Flag-WDR5B showed exclusive nuclear signals (Figure S1K) Although DCLK1 has been reported to bind microtubules, Flag-DCLK1 did not localize to primary cilia in RCTE cells (Figure S1F).^43^ It has been shown that EFHC2 exists on axonemes of motile cilia; however, we did not observe Flag-EFHC2 localizing to primary cilia (Figure S1G).^40^ Interestingly, Flag-CTDSP1 showed cilia localization but failed to co-localize with α-tubulin (Figure S1C and S1D). In contrast, three tagged proteins CCDC34, PSRC1, and C7orf57 localized to primary cilia and co-localized with cytoplasmic microtubules (Figure 3B-3G). PSRC1 was previously reported to be a microtubule binding protein, but its localization in primary cilia was unknown, whereas C7orf57 (renamed as FCAP33) was independently reported while this project was ongoing but the microtubule association has not been characterized.^44,45^ Together, our co-localization analyses identify CCDC34, PSRC1, and C7orf57/FCAP33 as three previously uncharacterized ciliary MAPs.

**Figure 3.**
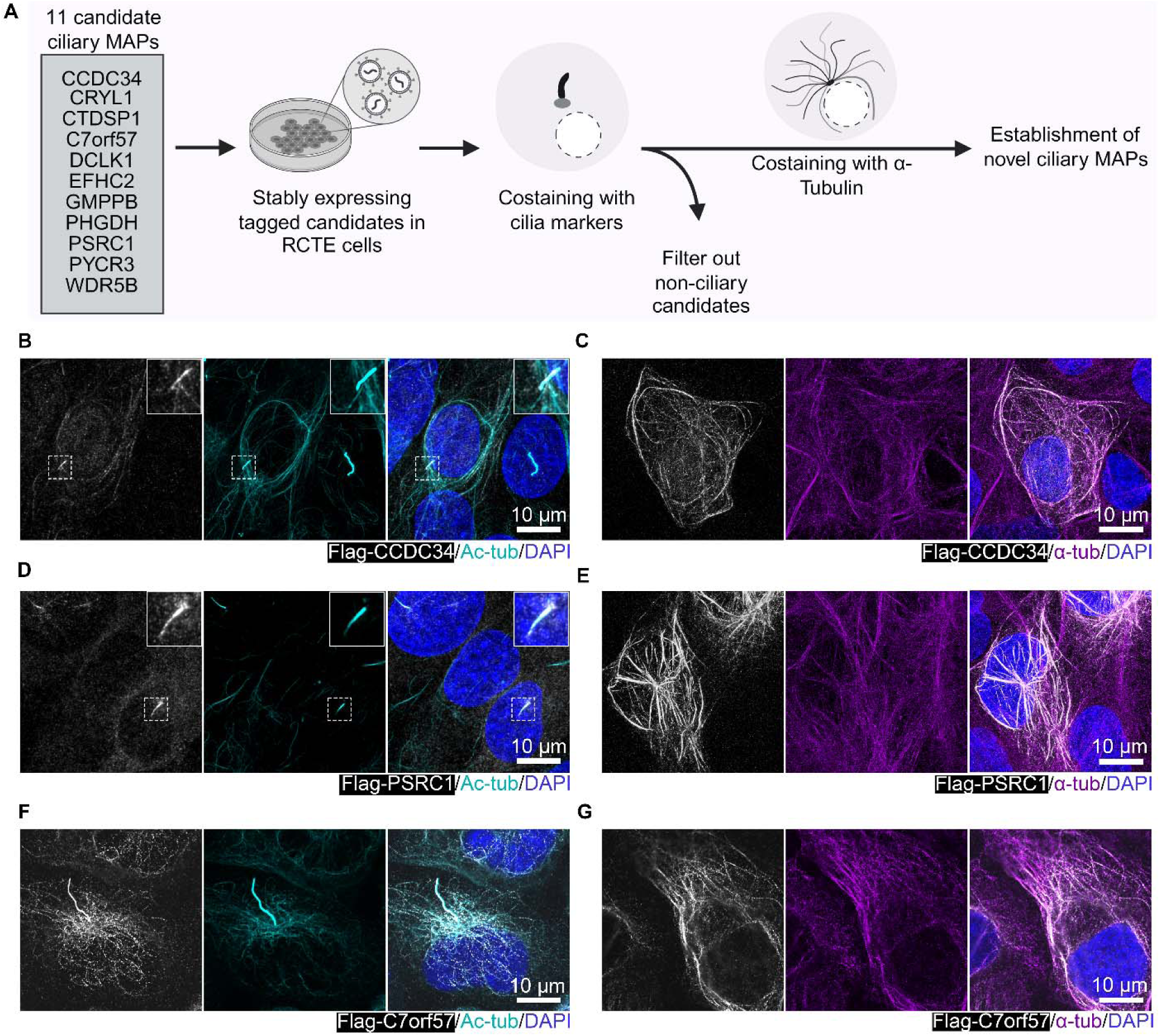
Candidate screening supports CCDC34, PSRC1, and C7orf57 as candidate ciliary MAPs. (A) Schematics showing co-localization studies of eleven candidate ciliary MAPs with cilia and microtubule markers to establish novel ciliary MAPs using a lentivirus mediated expression system in RCTE cells. Acetylated-tubulin (Ac-tub) is co-localized with (B) Flag-CCDC34, (D) Flag-PSRC1, and (F) Flag-C7orf57. α-tubulin (α-tub) is co-localized with (C) Flag-CCDC34, (E) Flag-PSRC1, and (G) Flag-C7orf57. All experiments were performed in triplicate.

### A poorly characterized Septin-9 isoform is enriched in the CEP41 proximity proteome

Bottom-up proteomics approach can provide isoform-specific peptide evidence, but shared peptides and database coverage limit isoform assignment.^46^ To identify isoform-associated enrichment in the CEP41 proximity proteome, we set the criteria that at least two unique peptides need to be assigned to one specific non-canonical isoform. Among candidates with isoform assignments, only the Septin-9 isoform, Septin-9_i8 (UniProt ID: Q9UHD8-8) met our criteria (Data S1 and Figure 4A and 4B). Specifically, two peptides were mapped to the unique N-terminus of Septin-9_i8, and one peptide spanned across the exon-exon junctions. Moreover, Septin-9 assigned by shared peptides is not found to be enriched in CEP41 proximity proteomes, indicating that this can be an isoform-biased enrichment (Figure 4C). Surprisingly, although being annotated in UniProt database, this isoform cannot be found in NCBI Refseq database. Literature search confirms that this isoform was once annotated as “NM_001113495.1” but then replaced by isoform f (NM_001113495.2). Together, these analyses provide peptide evidence consistent with enrichment of a poorly characterized Septin-9 isoform in the CEP41 proximity proteome.

**Figure 4.**
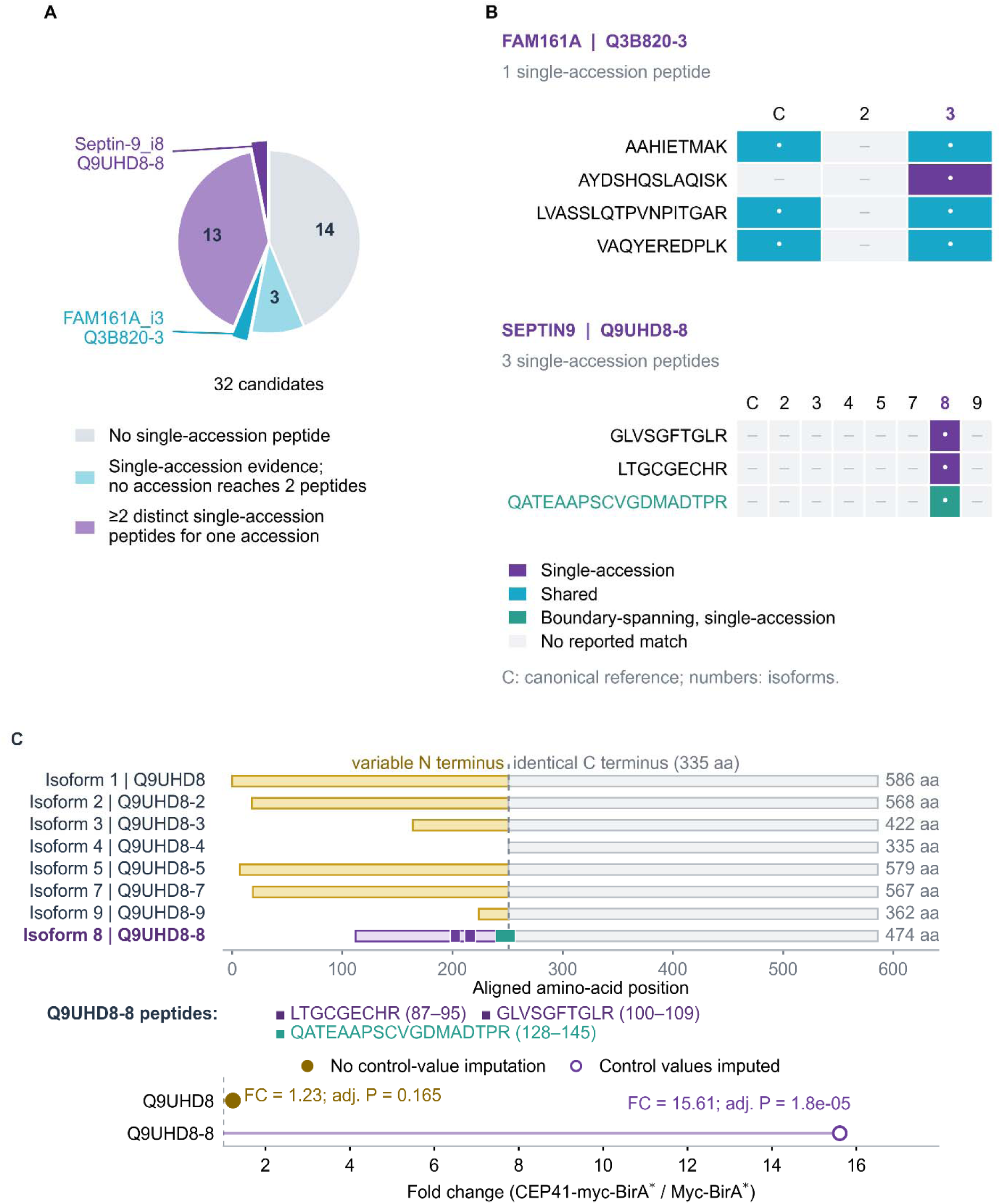
A poorly characterized Septin-9 isoform is enriched in the CEP41 proximity proteome. (A) Classification of 32 candidate proteins according to single-accession peptide evidence. 18 candidates have single-accession peptide evidence: noncanonical isoform-associated peptides are identified for FAM161A isoform 3 (Q3B820-3) and Septin-9 isoform 8 (Q9UHD8-8), highlighted by the exploded slices, while the other 16 candidates are assigned with canonical sequences. Purple slices represent candidates having at least two single-accession peptides. Cyan slices represent candidates having only one single-accession peptides. Gray slices represent candidates with no findings of single-accession peptides. (B) Peptide assignments for FAM161A and Septin-9. Rows show peptide sequences and columns show protein accessions. “C” represents canonical isoforms, and numbers represent noncanonical isoforms. Purple boxes indicate single-accession peptides, cyan boxes indicate shared peptides, and gray boxes indicate peptides not found in designated isoform sequences. The green box represents the boundary spanning peptide. (C) Comparison of reference Septin-9 isoform sequences and locations of the three peptides assigned to Q9UHD8-8. The lower plot shows the reported protein-group fold changes and adjusted p values for Q9UHD8 and Q9UHD8-8 in CEP41-myc-BirA* versus Myc-BirA* samples. Open and filled circles indicate groups with and without control-value imputation, respectively.

After excluding eight candidates as either non-ciliary proteins or non-ciliary MAPs and including a poorly characterized Septin-9 isoform, we retained a final list of high confidence CEP41 ciliary proximity proteome containing 32 primary cilia proteins (Data S2). To investigate potential function of these proteins, we designed two targeted screens on axoneme glutamylation and cilia architecture, respectively.

### Targeted RNAi screening does not identify robust regulators of axoneme glutamylation under the tested conditions

As CEP41 has been reported to regulate axoneme glutamylation in primary cilia, we asked whether CEP41 proximal proteins may also contribute to the regulation of axonemal glutamylation^21^ We selected hTERT-RPE1 cell line for this screen because of its relative uniform glutamylation staining in observations and the previously validated effects of CEP41 on axoneme glutamylation in this cell line.^47^ We excluded enzymes with established functions in tubulin PTMs and candidates whose roles in glutamylation had already been investigated. Because tubulin glutamylation occurs on the outer surface of microtubules, we did not prioritize Microtubule Inner Proteins (MIPs) for this targeted screen.^14^ Finally, we omitted proteins with low to no transcript levels in hTERT-RPE1 cells as those proteins are less likely to contribute to axonemal glutamylation in this cell line. The final list contains 19 candidates that were subjected to RNAi screen. (Data S2 and Figure S2A). RT-qPCR showed at least 70% transcript depletion for all tested siPools except those targeting TBC1D31 and HYLS1 (Figure S2B). We then measured relative axonemal GT335 fluorescence after depletion of each candidate. Most measurements fell between 80% and 120% of the control intensity. We therefore set over 20% change relative to a control as an exploratory effect-size threshold to prioritize candidates showing larger changes in axonemal polyglutamylation. Although depletion of BBS5 and SUFU results in statistically significant changes in GT335 intensity, the effect sizes were below the 20% threshold used for candidate prioritization.

Of note, the first Septin-9 siPool increased axoneme glutamylation level by more than 50%, whereas the second siPool did not reproduce this effect (Figure S2C). Thus, the targeted RNAi screen did not identify a candidate with a reproducible, robust effect on axoneme glutamylation.

### C7orf57 positively regulates primary cilium length

Defects in ciliopathy-associated proteins frequently alter cilia architecture. We therefore examined whether the newly identified ciliary MAPs affect cilia structure. According to the Human Protein Atlas, CCDC34 and PSRC1 are highly expressed in hTERT-RPE1 cells, while C7orf57 is expressed at low level in this cell type (Data S2). Given the expression profile of those three genes in hTERT-RPE1 cells, we performed RNAi-mediated knockdown on CCDC34 and PSRC1 and lentivirus-mediated overexpression on C7orf57 to determine their effects on cilia architecture (Figure 5A). We validated the knockdown efficiencies of PSRC1 and CCDC34 (Figure S2B and S3C). Depletion of either gene did not alter primary cilia formation rate and length (Figure S3A, S3B, S3D, and S3E). We also generated hTERT-RPE1 and RCTE cell lines stably expressing Flag-C7orf57 (Figure 5B). Compared with Flag control cells, Flag-C7orf57 overexpressing hTERT-RPE1 cells showed increased cilium length (3.04 ± 0.24 µm vs. 4.10 ± 0.12 µm), with no detectable change in the proportion of ciliated cells. (Figure 5C-5E). A similar increase in cilium length was also observed in RCTE cells, from 5.19 ± 0.20 µm to 7.51 ± 0.48 µm (Figure 5F-5H).

**Figure 5.**
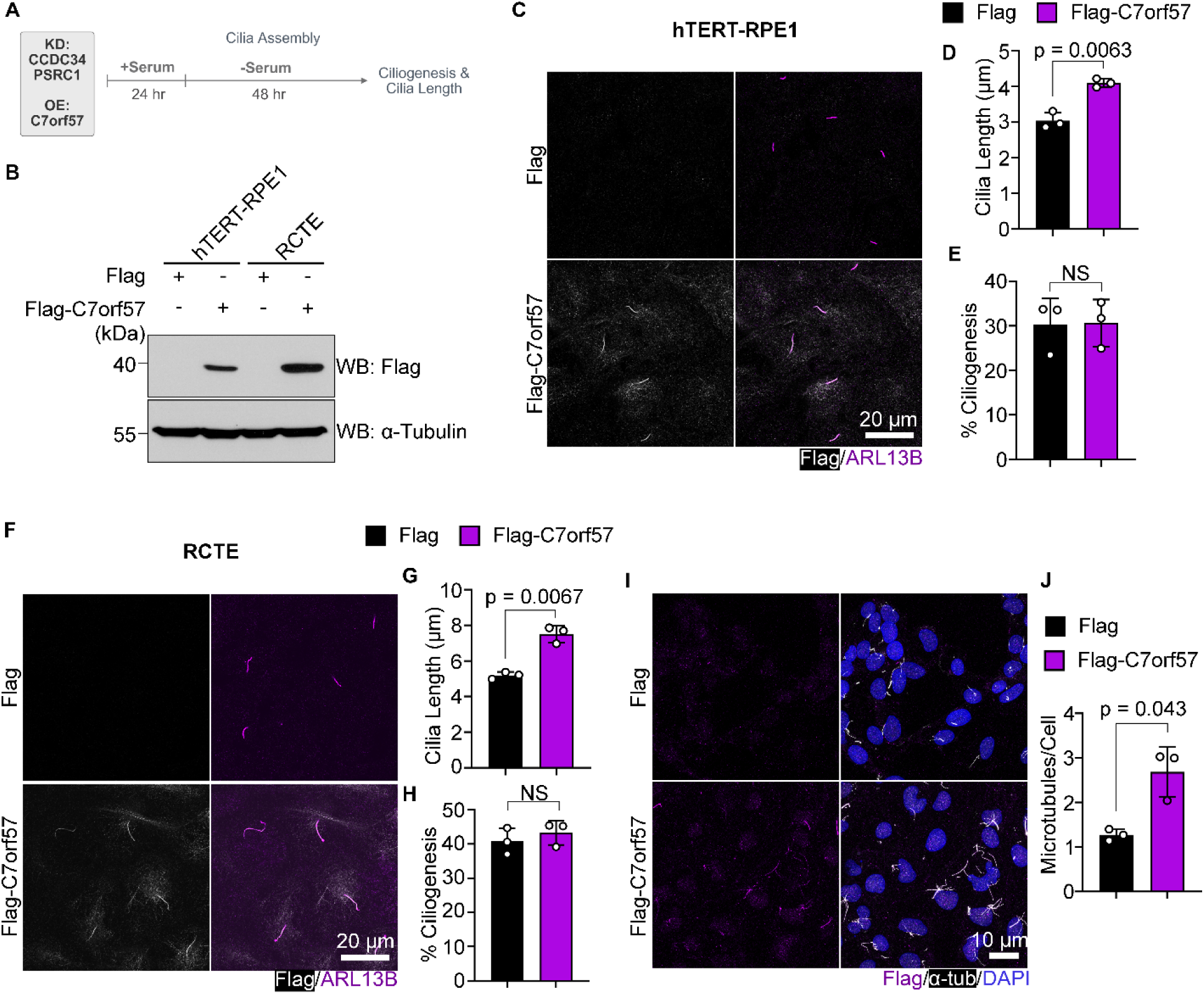
C7orf57 positively regulates primary cilium length. (A) Schematics of functional analysis of three newly identified ciliary MAPs on cilia architecture. (B) Validation of overexpressed Flag-C7orf57 in hTERT-RPE1 and RCTE cells. (C) Representative images and (D) quantification of cilia length in Flag and Flag-C7orf57 overexpressing hTERT-RPE1 cells. (E) Quantification of ciliogenesis rate in Flag and Flag-C7orf57 overexpressing hTERT-RPE1 cells. (F) Representative images and (G) quantification of cilia length in Flag and Flag-C7orf57 overexpressing RCTE cells. (H) Quantification of ciliogenesis rate in Flag and Flag-C7orf57 overexpressing RCTE cells. (I) Representative images and (J) quantification of residual microtubules in 5 µM nocodazole treated Flag and Flag-C7orf57 overexpressing RCTE cells. Significance was tested by two-tailed t-test with Welch’s correction. Error bars denote ±SD. All experiments were performed in triplicate.

Primary cilium length (PCL) control is regulated by multiple signaling pathways and biological processes.^48^ Because endogenous C7orf57 expresses at very low level in common human ciliated cell lines, we asked if stimuli that regulate PCL can transiently increase C7orf57 expression and subsequently contribute to stimuli-mediated PCL regulation (Figure S4A). We selected established and more recently discovered PCL regulating pathways and biological processes, including cAMP signaling, DYRK signaling, actin dynamics, calcineurin and GSK3β activities.^48–50^ We modulated these pathways and biological processes by treating hTERT-RPE1 cells with either inhibitors or activators. Neither short (3 or 6 hours) nor prolonged (24 hours) treatment produced a detectable increase in C7orf57 protein level, indicating that C7orf57-dependent PCL regulation is likely independent of these pathways (Figure S4B-S4C).

Only a handful of ciliary MAPs have previously been reported to regulate PCL when overexpressed. Dacheux and his colleagues discovered SAXO1, a MAP6-related family protein, regulates PCL by controlling the balance between axoneme assembly and disassembly. Specifically, SAXO1 was proposed to increase PCL through stabilizing axoneme microtubules and thus slowing disassembly rate. Importantly, MAP6 related family proteins can also promote resistance to drug- or cold-induced microtubule disassembly.^51^ In light of this model, we first examined whether C7orf57 overexpression altered the ciliary abundance of IFT components. Mean ciliary fluorescence of IFT88 and IFT140, components of IFT-B and IFT-A, respectively, was not detectably changed (Figure S4D and S4E). To determine if C7orf57 overexpression increases microtubule resistance to depolymerization, we treated RCTE cells with nocodazole and compared the residual microtubule networks. Interestingly, Flag-C7orf57 overexpressed cells contained more microtubules than Flag control cells, consistent with increased resistance to nocodazole-induced disassembly, resembling the phenotype reported for MAP6-related proteins (Figure 5I and 5J).

### C7orf57 is a paclitaxel-sensitive MAP that is associated with microtubules through its N-terminus

To map regions required for C7orf57 microtubule association, we analyzed its sequence and identified two predicted disordered regions (residues 33-52 and 180-295: NCBI NP_001093629.1). We then generated truncation variants that included or excluded those disordered regions (Figure 6A). Because MAPs frequently possess internal short repeats to facilitate their microtubule binding, we also searched for repeated sequence motifs.^52,53^ We performed a *de novo* identification of internal repeats in C7orf57 using HHrepID and MEME analysis, and identified three short tandem repeats, D-[W/Y]-[Y/M]-[I/V/Y]-H, in the N-terminal region of C7orf57.^54,55^ A more divergent DLLKH motif was identified only by HHrepID (Figure S5A and S5B). Because the four five-residue core motifs begin with aspartate and end with histidine, we termed then “DH repeats”. We generated a C7orf57 mutant construct lacking all four DH repeats (Figure 6A). Co-staining with α-tubulin showed microtubule association for the 1-179 and 53-295 variants, reduced association for the 53-179 variant, and no detectable association for the C-terminal 180-295 variant. The DH-repeat deletion failed to show detectable microtubule association (Figure 6B). These results implicate the importance of N-terminal region in C7orf57 microtubule association, although effects of the deletions on folding or stability remain possible.

**Figure 6.**
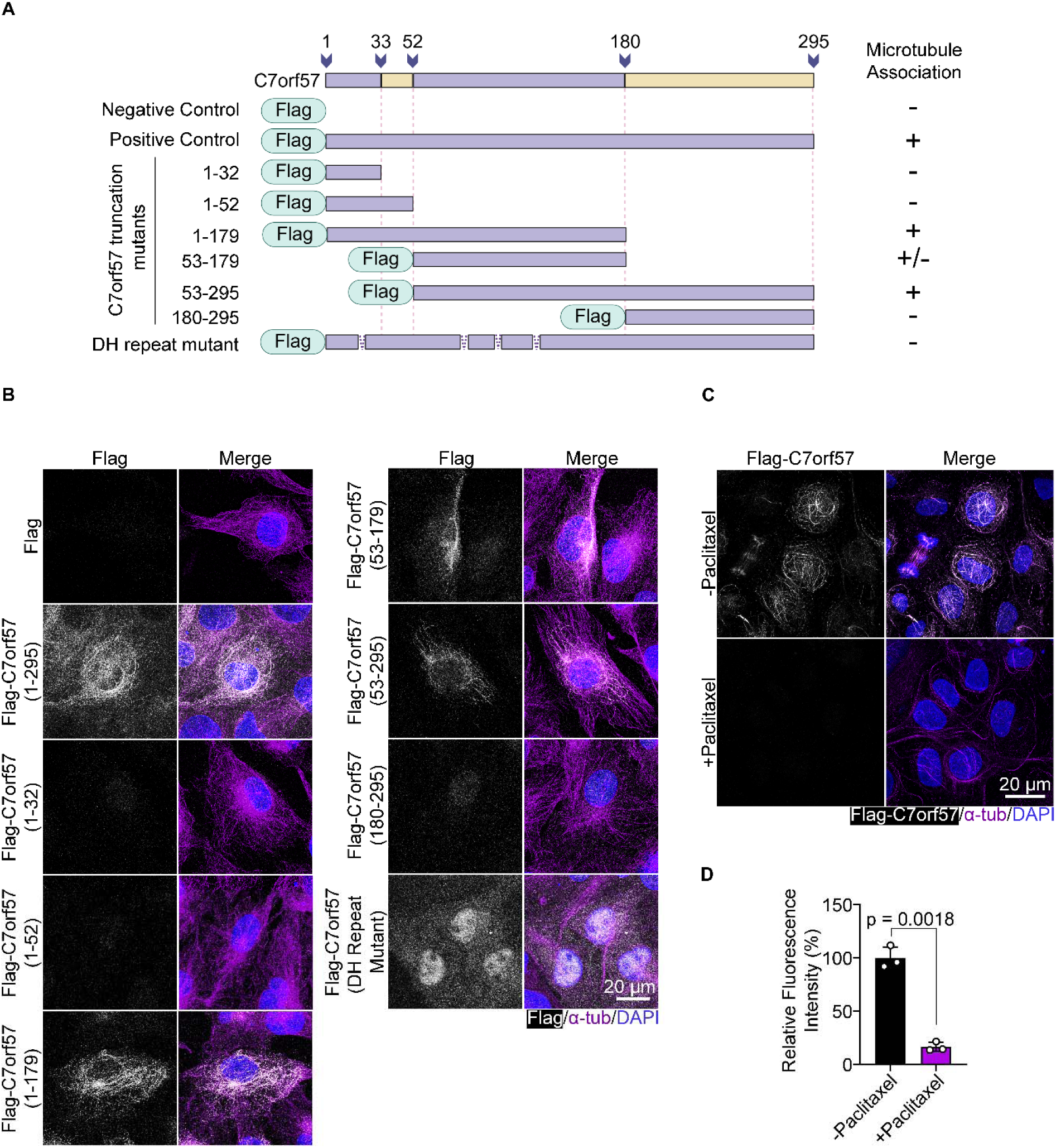
C7orf57 is a paclitaxel-sensitive MAP that is associated with microtubules through its N-terminus. (A) Schematics of full length and truncated Flag-tagged C7orf57. Yellow regions represent disordered regions predicted in NCBI (NP_001093629.1). Truncation mutants of C7orf57 were made based on those disordered regions. (B) Flag-tagged C7orf57 variants were co-stained with α-tubulin in RCTE cells to determine if those variants can still be detected on cystoplasmic microtubules. “+” represents positive microtubule association of C7orf57 variants. “-” represents negative microtubule association of C7orf57 variants. “+/−” represents reduced microtubule association of C7orf57 variants. (C) Paclitaxel reduces Flag-C7orf57 microtubule association. RCTE cells were treated with or without 500 nM paclitaxel for 12 hours before harvesting. (D) Quantification of relative fluorescence intensity of Flag-C7orf57 treated with or without paclitaxel. Significance was tested by two-tailed t-test with Welch’s correction. Error bars denote ±SD. All experiments were performed in triplicate.

Paclitaxel (Taxol) is a chemotherapy agent that stabilizes microtubules and perturbs mitosis.^56^ It binds to a site on β-tubulin facing the microtubule lumen, and can alter the association of MAPs with the lattice.^57–59^ To determine whether C7orf57 microtubule association was also affected by paclitaxel, we treated RCTE cells stably expressing Flag-C7orf57 with 500 nM paclitaxel for 12 hours. Flag-C7orf57 fluorescence intensity after paclitaxel treatment was markedly reduced compared to untreated RCTE cells (Figure 6C and 6D), furthering supporting the microtubule association feature of C7orf57.

## DISCUSSION

In this study, we used the ciliary MAP, CEP41, as a bait to define a cilia-enriched CEP41 proximal proteome and facilitate candidate ciliary MAP identifications. By integrating cilia-focused datasets with systematic filtering and manual curation, we generated a prioritized list of 32 CEP41 proximal cilia-associated proteins for downstream ciliary MAP identification. This prioritized dataset offers a focused starting point for investigating poorly characterized proteins associated with the ciliary microtubule compartment. Downstream screening identified three novel ciliary MAPs, CCDC34, PSRC1, and C7orf57. In addition, we confirmed an under-annotated Septin-9 isoform, denoted as Septin-9_i8, in our CEP41 proximity proteome. Among these candidates, functional analysis reveals C7orf57 as an important regulator of primary cilia length. Further studies in mammalian cells found that DH repeats in the N-terminal regions are crucial for C7orf57 microtubule association, and this association is sensitive to paclitaxel treatment.

During bait selection, we found that endogenous or low-level expressed exogenous CEP41 was exclusively detected at the axoneme and basal bodies, with little cytoplasmic microtubule localization. This selectivity raises an interesting question of how primary cilia recruit and retain distinct ciliary MAPs. One possibility is that ciliary MAPs recognize PTM patterns enriched along the axoneme. For instance, the microtubule binding of Tau protein, katanin, and KIF5B, is affected by tubulin polyglutamylation.^60^ In addition to glutamylation readers, a pool of MAPs also responds to tyrosination/detyrosination cycles of microtubules.^61^ Because axonemes are extensively modified with tubulin PTMs, MAPs with higher affinity towards microtubules bearing these PTMs may preferentially bind axoneme rather than dynamic cytoplasmic microtubules. Doublet geometry may provide a second layer of selectivity. Structural analysis of motile cilium doublets has shown that certain ciliary MAPs span across the junctions between A- and B-tubule.^62^ Compartment-specific scaffolds may provide a third mechanism. Like CEP41, low-level CSPP1 expression is restricted to centrosomes and centriolar satellites, while high-level expression produces cytoplasmic-microtubule bindings.^7,39^ Interestingly, depletion of PCM1, a pericentriolar scaffold protein, releases CSPP1 onto cytoplasmic microtubules.^39^ A comprehensive understanding of ciliary MAPs recruitments to primary cilia can help us better understand how axoneme homeostasis is maintained.

CCDC34 and PSRC1 are two novel ciliary MAPs identified in this study that showed no detectable effects on primary cilia architecture. Interestingly, deficiency of CCDC34 causes multiple morphological abnormalities of the sperm flagella (MMAF) with oligoasthenoteratozoospermia. Mechanistically, CCDC34 deficiency results in malformation of IFT-B complex in sperm flagella.^63^ However, we did not observe structural abnormalities of primary cilia in CCDC34-depleted hTERT-RPE1 cells, and this result likely indicates that IFT-B complex is not perturbed as well. One possible explanation for the phenotypic difference is functional redundancy in the primary cilium context. Related examples include overlapping functions of the ARPKD-associated protein DZIP1L with DZIP and ANKRD26 in ciliary regulation.^64,65^ It is therefore plausible that proteins functionally redundant with CCDC34 are expressed in primary cilia but not sperm flagella. PSRC1 is another MAP required for proper cell cycle at mitotic stages.^66^ PSRC1 has been functionally linked to CEP290 by binding and recruiting it to mitotic centrosome.^67^ Given CEP290 deficiency leads to a wide range of ciliopathies, it will be interesting to determine whether PSRC1 and CEP290 act together in primary cilia.

Alternative splicing in the MAP2/Tau family of MAPs represents a classic example of how various isoforms of MAPs are implicated in distinct physiological and pathological processes.^53^ Functional isoforms remain less well characterized in primary cilia. One example is Outer Dense Fiber protein 2 (ODF2), where different ODF2 isoforms occupy distinct ciliary compartments. In addition, only cenexin-1 (isoform 9 of ODF2) was able to regulate ciliogenesis.^68^ This raises the possibility that additional isoforms of ciliary proteins, including ciliary MAPs, have distinct roles in primary cilia. Indeed, Hansen and his colleagues observed that Septin-9 can be detected in primary cilia of hTERT-RPE1 and RPTEC but not ASC52telo cells.^69^ However, Septin-9 transcript level is comparable between ASC52telo and two other ciliated cell lines according to Human Protein Atlas.^70^ Because Septin-9 transcripts undergo alternative splicing, differential isoform expression could contribute to this heterogeneity. However, differences in protein abundance, localization, or antibody recognition are also possible. A comprehensive profiling of ciliary gene isoforms may improve our understanding of primary cilia composition and regulation.

Changes in primary cilium length may alter the abundance or local concentration of ciliary receptors and signaling components, with context-dependent effects on SHH and GPCR pathways.^48^ Defining the expression patterns of C7orf57 is therefore important for interpreting its potential physiological role. Although C7orf57/FCAP33 was independently identified in another primary cilium proteomic study, endogenous C7orf57 was below the immunoblot detection limit in the hTERT-RPE1, RCTE, and HEK293T cells tested here.^45^ This observation limits conclusions about endogenous C7orf57 functions and does not exclude enrichment of a small protein pool within cilia. Primary cilia proteome among tissues can show heterogeneity associated with their specialized functions.^71^ Ciliogenesis can also be induced in contexts such as senescence and skeletal-muscle differentiation.^72,73^ C7orf57 may therefore function in a restricted set of cell types, physiological conditions, or developmental stages. C7orf57 has also been reported in motile cilia.^74^ Studies of endogenous C7orf57 in cell types with stronger expression, together with *in vivo* phenotyping and human genetic analysis, will be important for defining its physiological and pathological relevance.^75,76^

Deletion of the DH repeats abolished detectable microtubule association of C7orf57. Such repeats contribute to microtubule association in several MAP families, including MDM1 and MAP2/Tau family proteins.^52,53^ Internal repeats can also mediate the microtubule-regulating activities of MAPs. For example, MAP6d1 Mn-motifs promote doublet microtubule assembly and tubulin recruitment.^77^ However, deletion of the DH repeats in C7orf57 may alter its folding or stability. Expression controls, targeted substitutions, and direct binding assays will be needed to define their contribution to the microtubule-binding interface. Paclitaxel markedly reduced the microtubule association of C7orf57. This effect is strikingly similar to that on a group of MIPs (i.e. JPT2, CSPP1 and JPT1), for which paclitaxel decreases association of MIPs with microtubules significantly by spatially limiting the intralumenal accessibility.^78^ This observation raises the possibility that C7orf57 may be a MIP. Nevertheless, paclitaxel can also alter the overall microtubule geometry and thus influence binding of outer surface MAPs, such as Doublecortin.^58^ Advanced imaging and structure biology techniques will be needed to determine whether C7orf57 is a bona fide MIP.

Besides newly discovered ciliary MAPs, the dataset generated in this study may have broader applications. As CEP41 itself is a Joubert syndrome (JBTS) and autism spectrum disorder (ASD) related protein, further characterization of CEP41-proximal proteins may advance our understanding of JBTS and ASD pathogenesis.^21,79^ Our dataset therefore provides a resource for prioritizing and identifying candidates that either physically or functionally interact with CEP41 in the context of ciliopathies. Our prioritization workflow was deliberately enriched for proteins already linked to cilia biology, which improves candidate selection but limits unbiased discovery, with the possibility of missing CEP41 proximal proteins that have dual roles in both cytoplasm and cilium. Additional ciliary MAPs may remain in the uncurated dataset. For example, γ-tubulin interacting protein, JPT1, is present in the original 365 significant proteins but not the final CEP41 proximal proteome list.^80^ LGALSL is another recently identified MAP that is also present in the uncurated dataset.^81^ It will be interesting to determine whether those additional MAPs may function in primary cilia. In conclusion, our proof-of-principle studies support the use of this dataset as a resource for future identification and characterization of ciliary MAPs and candidate ciliopathy-associated proteins.

### Limitations of the study

Several limitations constrain interpretation. First, ciliary proteomes show high diversity across tissues. Primary cilia in different parts of the body may recruit unique ciliary MAPs to support their functions.^71^ The CEP41 proximity proteome was generated in RCTE cells, whereas selected candidates were also evaluated in hTERT-RPE1 cells; the results may therefore not generalize to other tissues. Second, residual cytoplasmic biotinylation and expression of the fusion bait can contribute non-ciliary proteins to the dataset. Third, manual curation enriched for proteins already associated with cilia and may have excluded unexpected candidates. Fourth, BioID and co-localization do not establish direct binding, and overexpression phenotypes do not demonstrate an endogenous requirement for C7orf57. Finally, the primary cilium is highly compartmentalized, with distinct cellular processes, such as ectocytosis and ciliary gating, occurring at specialized subciliary regions^1^. A broadly distributed bait (e.g. CEP41) may therefore underrepresent low-abundance proteins at the ciliary tip, transition zone, or other subciliary compartments. Future studies should combine orthogonal validation with compartment-specific profiling across ciliated cell types.

## RESOURCE AVAILABILITY

### Lead contact

Requests for further information and resources should be directed to and will be fulfilled by the lead contact, Jinghua Hu.

### Materials availability

All unique/stable reagents generated in this study are available from lead contact upon reasonable request.

### Data and code availability

- The mass spectrometry data generated in this study have been deposited to the ProteomeXchange Consortium under the dataset identifier PXD085758 and is publicly available as of the date of publication.
- Original western blot images and data points for quantification are included in Data S3. Microscopy data reported in this paper will be shared by the lead contact upon request.
- All original code has been deposited at GitHub (https://github.com/Yan-Liang-2023/CEP41-BioID) and is publicly available as of the date of publication.
- Any additional information required to reanalyze the data reported in this paper is available from the lead contact upon request.

## ACKNOWLEDGMENTS

We would like to thank the Proteomics and Metabolomics Facility at The Wistar Institute for providing proteomics services. This study was supported by Mayo Clinic Graduate School of Biomedical Sciences, the Sigma Xi Grants in Aid of Research program and National Institutes of Health (Project number: R01-DK099160; R01-AG076469).

## AUTHOR CONTRIBUTIONS

Conceptualization, X.S. and J.H.; methodology, X.S., Y.L. and J.H.; Investigation, X.S., Y.L. and J.H.; writing – original draft, X.S., Y.L., and J.H.; writing – review & editing, X.S., Y.L. and J.H.; funding acquisition, X.S. and J.H.; resources, J.H.; supervision, J.H.

## DECLARATION OF GENERATIVE AI AND AI-ASSISTED TECHNOLOGIES IN THE WRITING PROCESS

The authors carefully used LLMs during the preparation of this manuscript. After using these tools, the authors reviewed and edited the content as needed and took full responsibility for the content of the publication.

## SUPPLEMENTAL INFORMATION

Document S1. Figures S1–S5, Table S1 and S2

Data S1.

Data S2.

Data S3.

## METHODS

### KEY RESOURCES TABLE

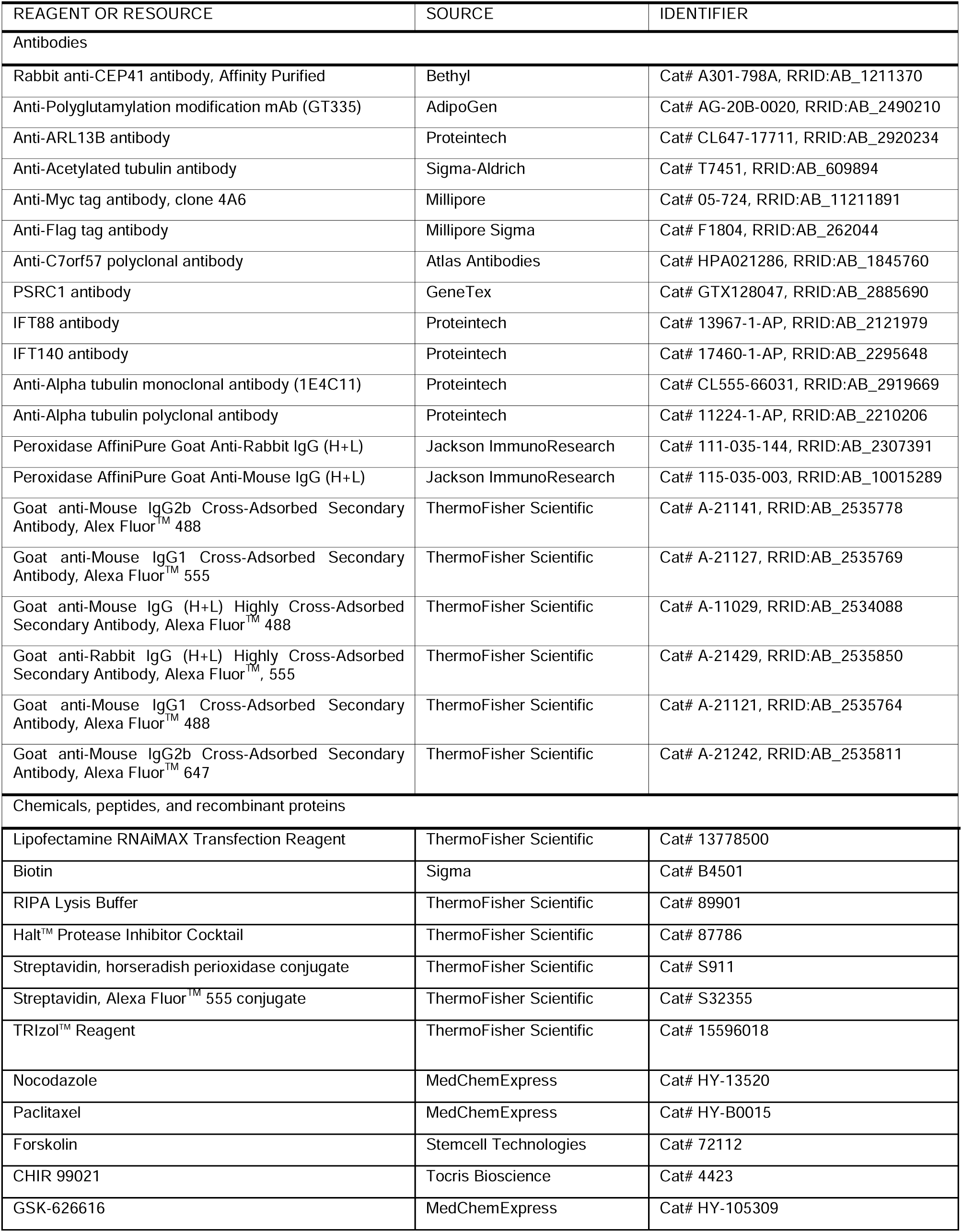
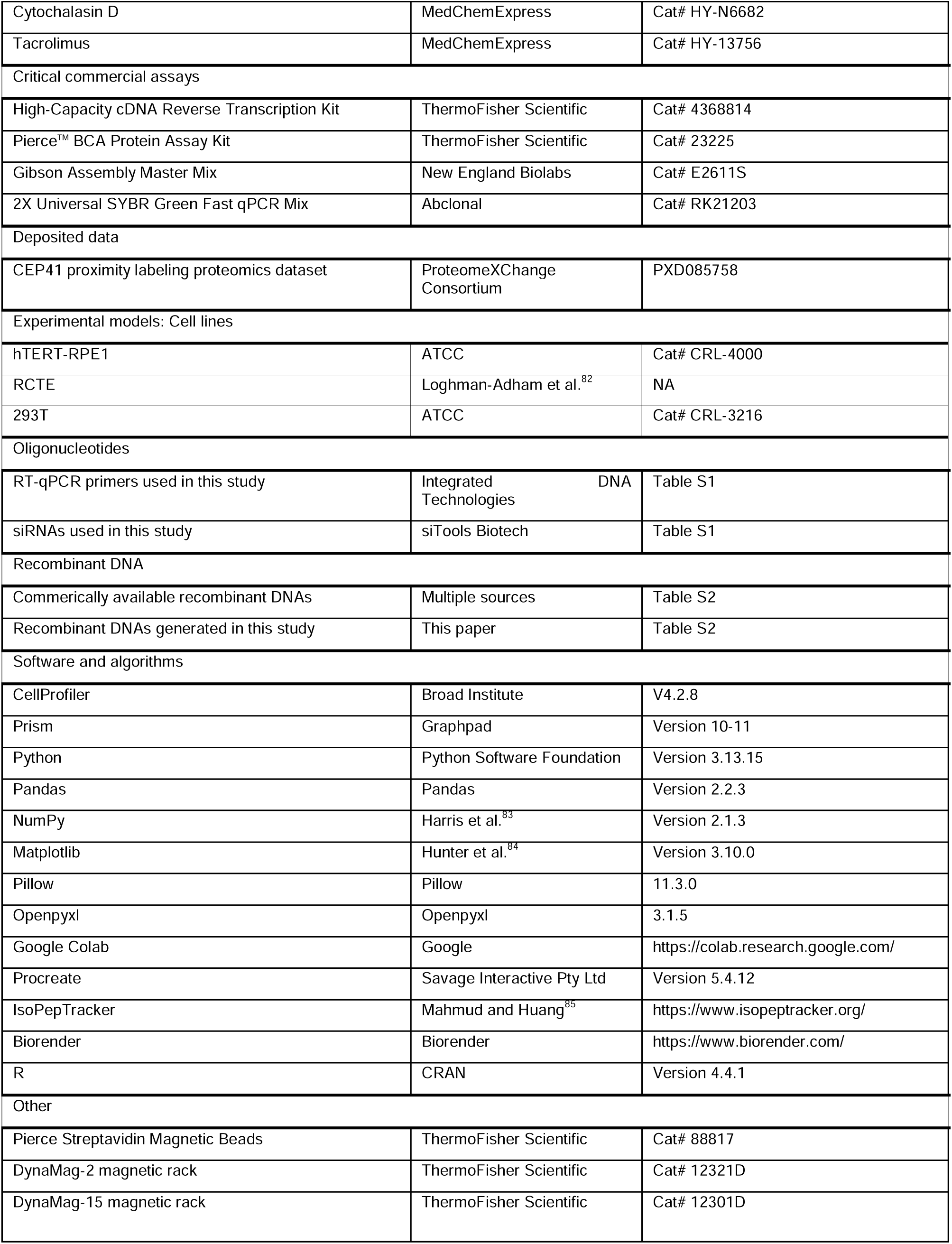

### EXPERIMENTAL MODELS AND STUDY PARTICIPANTS DETAILS

HEK293T cells (ATCC, CRL-3216) were cultured in Dulbecco’s Modified Eagle’s Medium (DMEM; Corning) supplemented with 10% Fetal Bovine Serum and 1% Pen/Strep at 37°C with 5% CO_2_. hTERT-RPE1 and Renal Cortical Tubular Epithelial (RCTE) cells were cultured in Dulbecco’s Modified Eagle Medium/Nutrient Mixture F-12 (DMEM/F-12; Gibco) supplemented with 10% Fetal Bovine Serum and 1% Pen/Strep at 37°C with 5% CO_2_.

### METHOD DETAILS

#### Molecular Cloning

All Plasmids except the C7orf57 DH repeat mutant were constructed using restriction endonuclease digestion and subsequent ligation to clone target fusion genes into our destination lentiviral vectors. For constructing plasmid bearing C7orf57 DH repeat mutant, we used Gibson Assembly to join five C7orf57 fragments that skip DH repeats in between.

#### Immunofluorescence staining and imaging

Cells were cultured on glass coverslips, rinsed with PBS before fixing with 4% paraformaldehyde (PFA) for 15 min at room temperature and washed three times with PBS. PBS with 0.1% (v/v) Triton-X100 was added for 10 minutes for permeabilization and washed three times with PBS before blocking with 3% BSA in PBS for 1 hour at room temperature. Primary antibodies diluted in blocking buffer were applied overnight at 4 °C, followed by PBS washes and fluorescent secondary antibodies incubation for 1 hour at room temperature. Confocal imaging with or without Z-stack was performed on Zeiss LSM 980 microscope with a 63x oil objective at 1.7x digital zoom. Laser channels at excitation wavelength of 405/488/555/647 nm were used depending on the experimental purposes. Zen microscopy software was used for orthogonal and Airyscan processing.

#### Immunoblot

Cells were lysed in RIPA buffer, and protein concentrations were determined using a Bicinchoninic acid assay (BCA) according to the kit instructions. Equal amounts of protein (30 µg per sample) were separated by gel electrophoresis (SDS-PAGE) and subsequently transferred onto membranes. Membranes were blocked with 5% non-fat milk for 1 hour at room temperature, and incubated with the primary antibodies overnight at 4 °C. The appropriate HRP-conjugated secondary antibodies were added for incubation 1 hour at room temperature on the following day. Protein signals were visualized using Enhanced Chemiluminescence (ECL). Membranes were stripped and re-probed with additional antibodies where experiments were required.

#### BioID and LC-MS/MS

##### BioID sample preparation

Cells were grown to confluence and serum-starved for 24 h, then incubated for an additional 24 h in starvation medium supplemented with 50 μM biotin. Following a PBS wash, cells were lysed in RIPA buffer containing 1× Halt Protease Inhibitor Cocktail. Samples were sonicated, clarified by centrifugation, and quantified. Equal amounts of protein were incubated with magnetic streptavidin beads overnight at 4°C. Beads were washed twice with lysis buffer, once with 1 M KCl, once with 0.1 M Na₂CO₃, once with 2 M urea in 10 mM Tris-HCl (pH 8.0), and twice with lysis buffer. Five percent of the bead volume was reserved for pulldown validation. The remaining samples were washed with 50 mM Tris-HCl (pH 7.5) followed by 2 M urea in 50 mM Tris-HCl (pH 7.5). After removal of the final wash, protein-bound beads were submitted to the Proteomics and Metabolomics Facility Shared Resource at The Wistar Institute for sample processing and LC-MS/MS analysis.

##### Mass Spectrometry

Six streptavidin bead-bound samples, triplicates of Myc-BirA* and hCEP41-myc-BirA*, were reduced with TCEP, alkylated with iodoacetamide, and digested on-bead with trypsin. The digests were desalted using centrifugal C18 columns (Nest Group). Digests were resuspended in 0.015% dodecyl maltoside, 0.1% trifluoroacetic acid, and 10% of each sample was injected for each sample.

LC-MS/MS analysis was performed using an Orbitrap Astral mass spectrometer (Thermo Fisher Scientific) coupled to a Vanquish Neo UPLC system (Thermo Fisher Scientific). For the chromatographic separation, the column compartment temperature was set to 45.0 °C, and the flow rate was set to 0.300 µL/min. Buffer A consisted of 0.1% formic acid in Milli-Q water and buffer B consisted of 0.1% formic acid in acetonitrile. Samples were loaded onto a trap column (nanoEase M/Z Symmetry C18, 100 Å, 120 μm x 20 mm, Waters), which was placed in line with an analytical column (Acclaim PepMap, 100 Å, 75 μm x 250 mm, Thermo), and the sample was separated using a 40 min linear gradient increasing from 5% to 30% B over 25 min, from 30% to 40% B over 6 min, from 40% to 80% B over 2 min, and held at 80% B for 7 min before re-equilibration.

MS data were acquired in data-independent acquisition mode. An MS^1^ scan was collected every 0.6 s in the Orbitrap at 240 k resolution. Ions were injected for 3 ms or until the 5e6 AGC target was reached. Precursor ions within a mass range of 380-980 *m/z* were collected in 2 *m/z* isolation windows with a maximum injection time of 3 ms or until an AGC target of 5e4 ions was reached. Precursors were fragmented using 25% normalized collisional energy. MS^2^ scans of fragments ranging from 110 – 2000 *m/z* were collected in the Astral mass analyzer.

##### Data Analysis

Raw mass spectrometry files were searched using DIA-NN v2.0.0^86^ against a spectral library generated from the SwissProt human database (UP000005640; downloaded August 6, 2024), supplemented with construct sequences and a contaminant database. N-terminal acetylation, N-terminal methionine excision, STY phosphorylation, and methionine oxidation were specified as variable modifications, while cysteine carbamidomethylation was set as a fixed modification. Searches were performed allowing one missed cleavage, up to two variable modifications per peptide, peptide lengths of 7-50 amino acids, and precursor charge states of 1-4. MS1 and MS2 mass tolerances were set to 8 and 15 ppm, respectively. Match-between-runs was enabled, and cross-run normalization was disabled.

Contaminants, proteins identified by a single peptide, and proteins with less than 3 valid values were filtered from the dataset. Protein abundance values were normalized to the median Myc-BirA* abundance. Differential expression analysis was performed using LFQ Analyst.^87^ Missing values were imputed with the minimum value of the filtered dataset, and FDR levels were controlled using the Benjamini-Hochberg correction. Proteins with a fold change of >2 and an adjusted p. value of <0.05 were considered significant.

#### Prioritization workflow for CEP41 proximity proteome

Significantly enriched proteins in samples of hCEP41-myc-BirA* over Myc-BirA* were overlapped with an inclusive cilia dataset.^25^ The resulting 68 candidates were subjected to a filter, which excludes tubulins, membrane-associated, non-ciliary, and mitochondrial proteins. Literature search was conducted to find experimentally validated non-ciliary proteins and membrane-associated proteins with references. MitoCarta 3.0 database was used to search mitochondrial proteins in our dataset.^88^ Filtering details are described in “Filter” sheet of Data S2. After removing unwanted candidates, we obtained a list of 40 candidates that can be found in “Curated List” sheet of Data S2. Further determination of 8 non-ciliary or non-ciliary MAP proteins generated the final list of 32 highly confident CEP41 primary ciliary proximity proteins that are described in detail in “Final List” sheet of Data S2. Proteins included in “Curated List” and “Final List” are annotated with 1) ciliary localization, ciliopathy association, microtubule association, transcript levels from Human Protein Atlas database, and functions in glutamylation with references.^70^

#### Isoform-associated peptide assignment and Septin-9 sequence analysis

Isoform-associated peptide evidence was evaluated using the “All Peptides” table sheet from the original DIA-NN output file (Data S1). The Protein Group field was used to identify the protein groups whereas the Protein Ids field was used to determine the individual protein accessions matched by each peptide. Peptide entries were grouped by stripped amnio-acid sequence so that different precursor charge states or modified forms of the same peptide were counted only once. For stripped sequence represented by multiple precursor or modified-sequence entries, the union of accessions reported across the corresponding Protein Ids fields was retained. A peptide was classified as a single-accession peptide when it was assigned to only one isoform. A peptide was considered isoform-informative when it matched some, but not all, protein isoforms of the corresponding gene represented in the original DIA-NN report. These classifications were defined relative to the protein-sequence database used for the original search.

Q9UHD8-8 was selected for further analysis because three experimentally detected peptides were assigned exclusively to this accession in the original DIA-NN report. To investigate their possible transcript context, the archived RefSeq SEPTIN9 transcript, NM_001113495.1, from NCBI database was analyzed using IsoPepTracker to predict its open reading frame and compare the translated protein sequence with reference Septin-9 isoforms.^85^

#### RNAi in mammalian cells

siPools were used to treat hTERT-RPE1 cells according to manufacturer’s instructions (siTools Biotech). Briefly, the mixture of 1 nM siPool and RNAiMAX diluted in Opti-MEM medium with reduced serum (Gibco) was added to cell culture. After 24 hours, cells were starved for another 48 hours to induce cilia formation. Cells were harvested for downstream assays, including western blot, RT-qPCR, and immunofluorescence.

#### Assays with drug treatments

##### Nocodazole treatment

RCTE cells were seeded on glass coverslips and cultured in complete media for 24 hours, then serum-starved for 48 hours to induce ciliogenesis. 5 µM nocodazole was added to the treatment group 1 hour prior to fixation. Cells were rinsed twice with PHEM buffer (60 mM PIPES, 25 mM HEPES, 4 mM MgSO_4_, 10 mM EGTA, pH7.0), then briefly extracted for 1 min with PHEM containing 0.2% Triton X-100 to remove soluble cytoplasmic components. Coverslips were gently washed with PBS and fixed as described above in immunofluorescence staining and imaging section.

##### Paclitaxel treatment

500 nM Paclitaxel was added 12 hours before fixation. Subsequent PHEM wash, detergent extraction, PBS rinse, and fixation steps were performed identically to the nocodazole-treated experiment.

##### PCL-regulating drugs treatment

hTERT-RPE1 cells were seeded and cultured until approximately 80% confluence. On the following day, cells were serum-starved for a total of 48 hours to induce ciliogenesis. During the starvation period, cells were treated with a panel of compounds previously reported to increase cilia length, including Forskolin (10 µM), CHIR99021 (5 µM), GSK-626616 (1 µM), cytochalasin D (1 µM), and FK506 (tacrolimus; 2.5 µM). DMSO-treated and untreated cells were included as vehicle and baseline control, respectively. Cells were treated with drug for 3, 6, or 24 hours. All treatment groups were collected at the end of the 48-hour starvation period.

#### Generation of cell lines with stably expressed constructs

To generate hTERT-RPE1 and RCTE cells stably expressing fusion proteins, we first transfected 293T cells with pCDH vectors containing target fusion genes, along with psPAX2 and PMD2.G plasmids. Then, HEK293T supernatants were collected and added to either hTERT-RPE1 or RCTE cells in the presence of 10 µg/ml polybrene. After 48 hours, cells were selected with Geneticin (800 µg/ml) for another week to eliminate cells without expressing constructs.

#### RT-qPCR

RNAs from cells were extracted using TRIzol reagent according to manufacturer’s protocols (ThermoFisher). Afterwards, extracted RNAs were reverse transcribed to cDNA using High-Capacity cDNA Reverse Transcription kit (ThermoFisher Scientific). Diluted cDNAs were then used as templates for subsequent RT-qPCR experiments using self-designed primers.

#### De novo identification of internal repeats in C7orf57 primary sequence

For HHrepID analysis, C7orf57 protein sequence (NP_001093629.1) was used as the input. Default settings were used for identification of novel internal repeats. The resulting repeats with p-values larger than 0.05 were omitted for downstream analysis.

For MEME analysis, the same input was used, and the default settings were again implemented. Discovered motifs with E-value less than 0.05 were selected and compared with results generated in HHrepID analysis.

### QUANTIFICATION AND STATISTICAL ANALYSIS

#### Quantification of assays with drug treatments

##### Nocodazole treatment

Cells were imaged using a Zeiss LSM 980 microscope with 63x oil objective without airyscan processing at 1x digital zoom. Imaging parameters were kept consistent across experimental groups. For each replicate, five randomly selected fields were manually quantified. Three independent replicates were analyzed for each experimental group.

##### Paclitaxel treatment

Cells were stained with DAPI, anti-α-tubulin, and anti-Flag (Flag-C7orf57), and imaged as RGB composite TIFF files per field. Automated quantification was performed in CellProfiler (v4.2.8; Broad Institute). Images were grouped by condition (control vs. Paclitaxel) and replicate (R1, R2 and R3) using metadata extracted from filenames. Each RGB composite was split into its red, green, and blue channels using the ColorToGray module, corresponding to Flag, α-tubulin, and DAPI, respectively. Nuclei were identified from the DAPI channel using the IdentifyPrimaryObjects module (diameter 85 - 200 pixel units). Cell boundaries were defined from the α-tubulin channel using the IdentifySecondaryObjects module with a watershed-gradient algorithm seeded from the identified nuclei. Total and mean Flag immunofluorescence intensity was measured per cell by the module MeasureObjectIntensity. Per-cell measurements, including object identity, condition, and replicate metadata, were exported as a single CSV file via the ExportToSpreadsheet Module for downstream statistical analysis.

#### Statistical Analysis

All statistical analyses were performed in GraphPad Prism. Each datapoint for quantification of primary cilia related experiments is an average of at least 50 primary cilia measurements. Means of triplicate were used to perform either two-tailed t-test with Welch’s correction or one sample t-test against hypothetical value of 100 (specifically for experiments shown in Figure S2). Data are presented in means ± SD. P-values less than 0.05 are considered statistically significant.

