## Supplemental Information for "Identification of candidate ciliary microtubule-associated proteins using CEP41 proximity labeling"

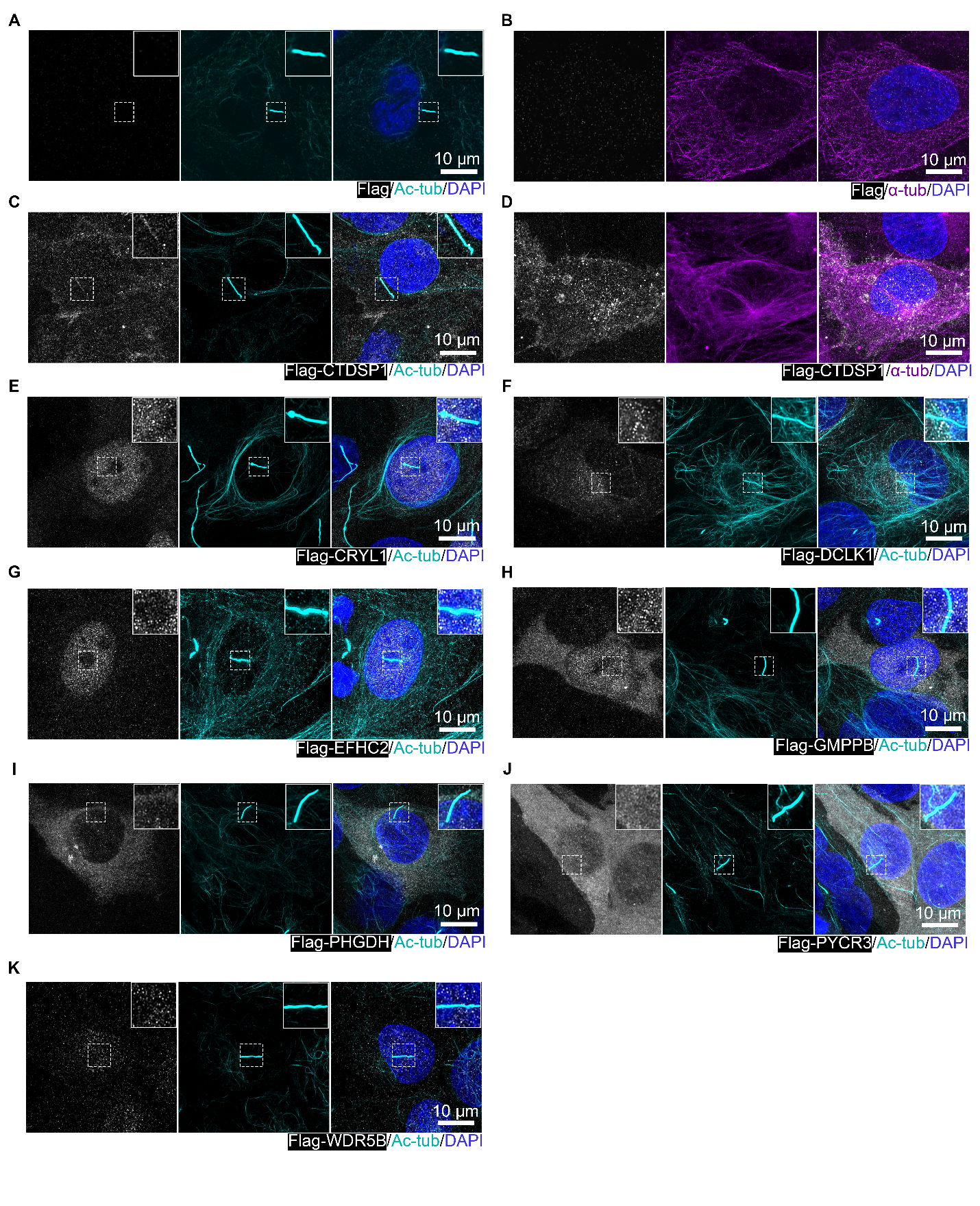
**Figure S1: Co-localization studies exclude eight candidates to be ciliary MAPs.** Acetylated tubulin is co-stained with (A) Flag, (C) Flag-CTDSP1, (E) Flag-CRYL1, (F) Flag-DCLK1, (G) Flag-EFHC2, (H) Flag-GMPPB, (I) Flag-PHGDH, (J) Flag-PYCR3, and (K) Flag-WDR5B. α-tubulin is co-stained with (B) Flag and (D) Flag-CTDSP1. n = 3 for both technical and biological replicates.

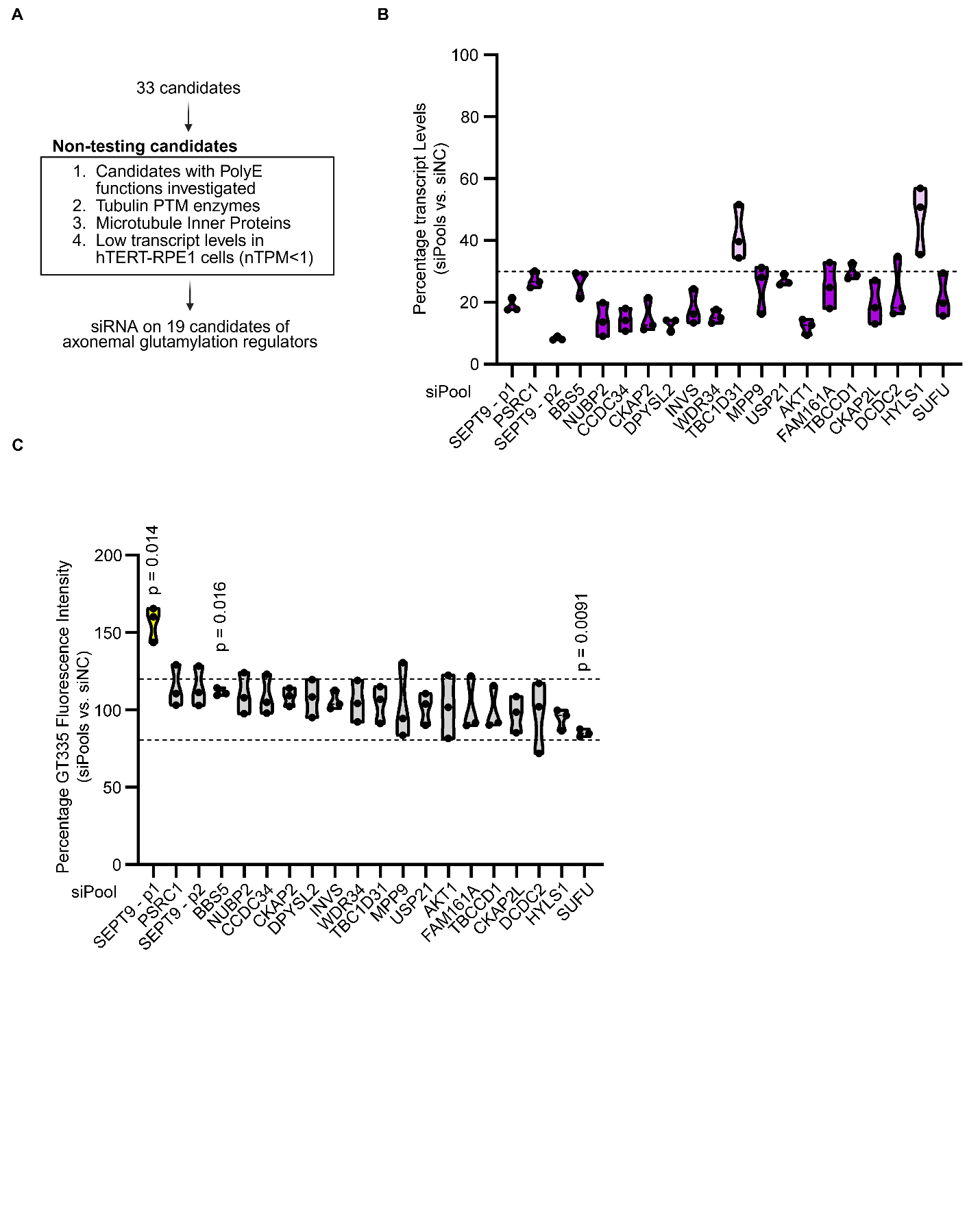
**Figure S2: Targeted RNAi screening does not identify robust regulators of axoneme glutamylation under the tested conditions.** (A) Schematics showing selection of 19 candidates for siRNA screen on identifying novel players in mediating axoneme glutamylation. (B) Validation of siPools’ knockdown efficiencies using RT-qPCR. Dash line represents 30% of transcript level in siPool treated cells as compared to control. (C) Targeted siRNA screen on 19 candidates. Dash lines represent subjective cutoff lines with lower limit being 80% and upper limit being 120% of axoneme GT335 fluorescence intensity in control cells. One sample t-test was performed against the hypothetical value of 100. Error bars denote ±SD. All experiments were performed in triplicate.

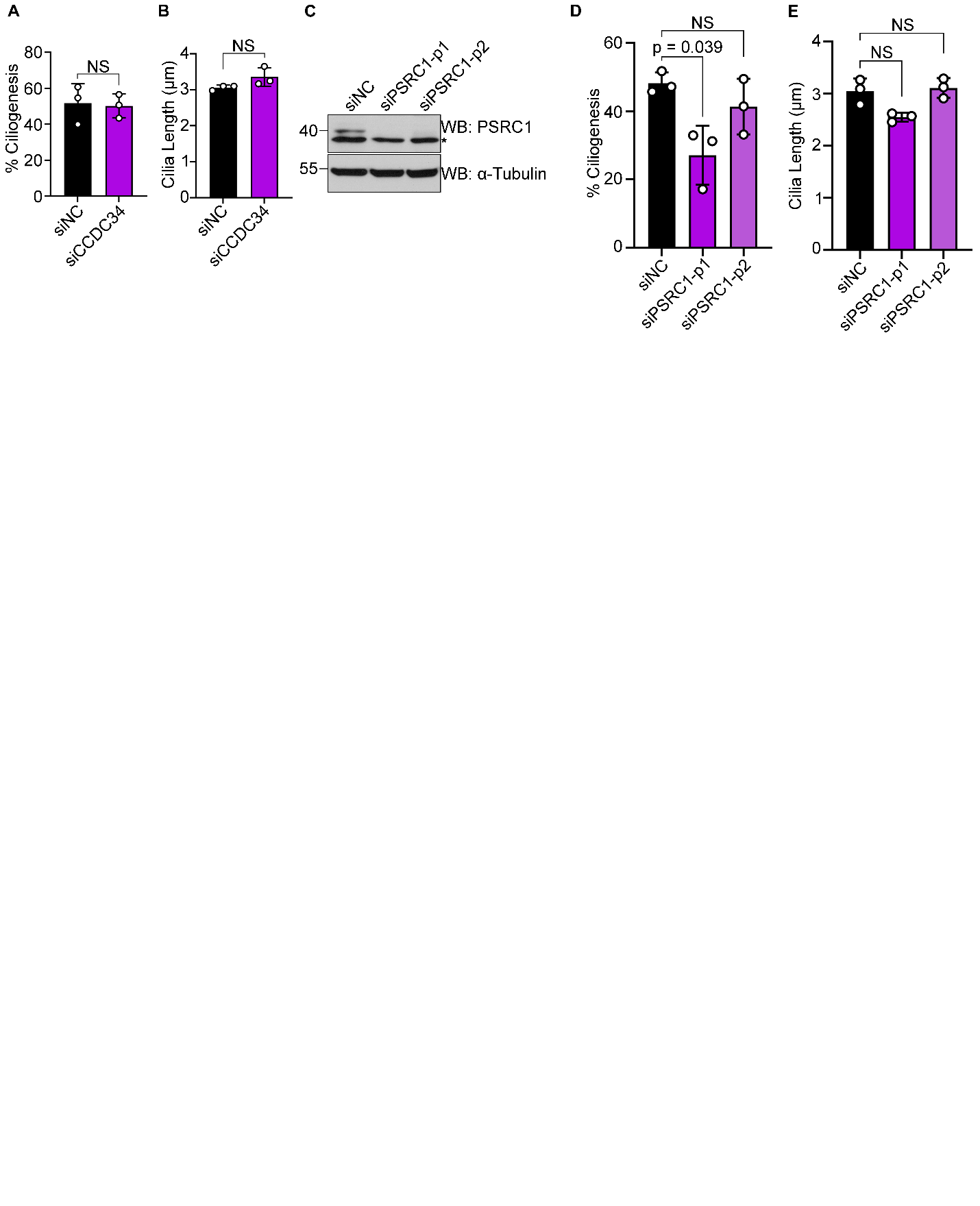
**Figure S3: CCDC34 and PSRC1 do not affect cilia architecture.** Knockdown CCDC34 does not affect (A) cilia formation and (B) cilia length. (C) Western blot validation of PSRC1 siRNA efficiencies. Asterisk represents non-specific bands. (D) Knockdown PSRC1 with siPSRC1-p1 but not siPSRC1-p2 results in decreased cilia formation rate. (E) PSRC1 siRNAs do not affect cilia length. Significance was tested by two-tailed t-test with Welch’s correction. Error bars denote ±SD. All experiments were performed in triplicate.

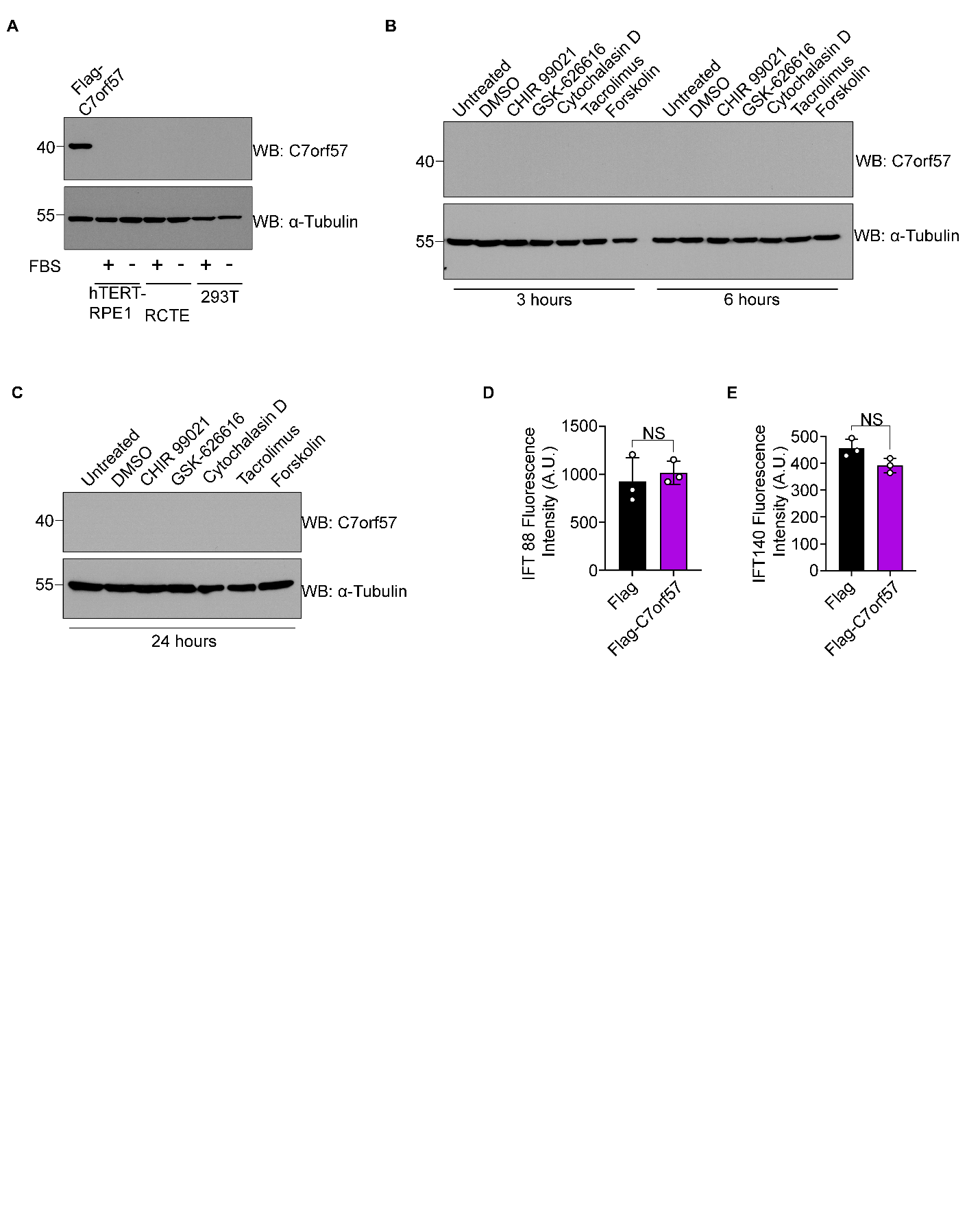
**Figure S4: Regulation of primary cilia length (PCL) by C7orf57 is independent of well-established cilia length regulating pathways and biological processes.** (A) C7orf57 expresses lowly in RPE, RCTE, and 293T with or without serum starvation. The sample from Flag-C7orf57 stably expressing hTERT-RPE1 cells serves as positive control. PCL-controlling drugs cannot induce transient upregulation of C7orf57 expression (B) 3 hours, 6 hours, or (C) 24 hours after treatments. C7orf57 overexpression does not change (D) anterograde and (E) retrograde Intraflagellar Transport (IFT) machinery represented by IFT88 and IFT140 mean ciliary intensity respectively in hTERT-RPE1 cells. Significance was tested by two-tailed t-test with Welch’s correction. Error bars denote ±SD. All experiments were performed in triplicate.

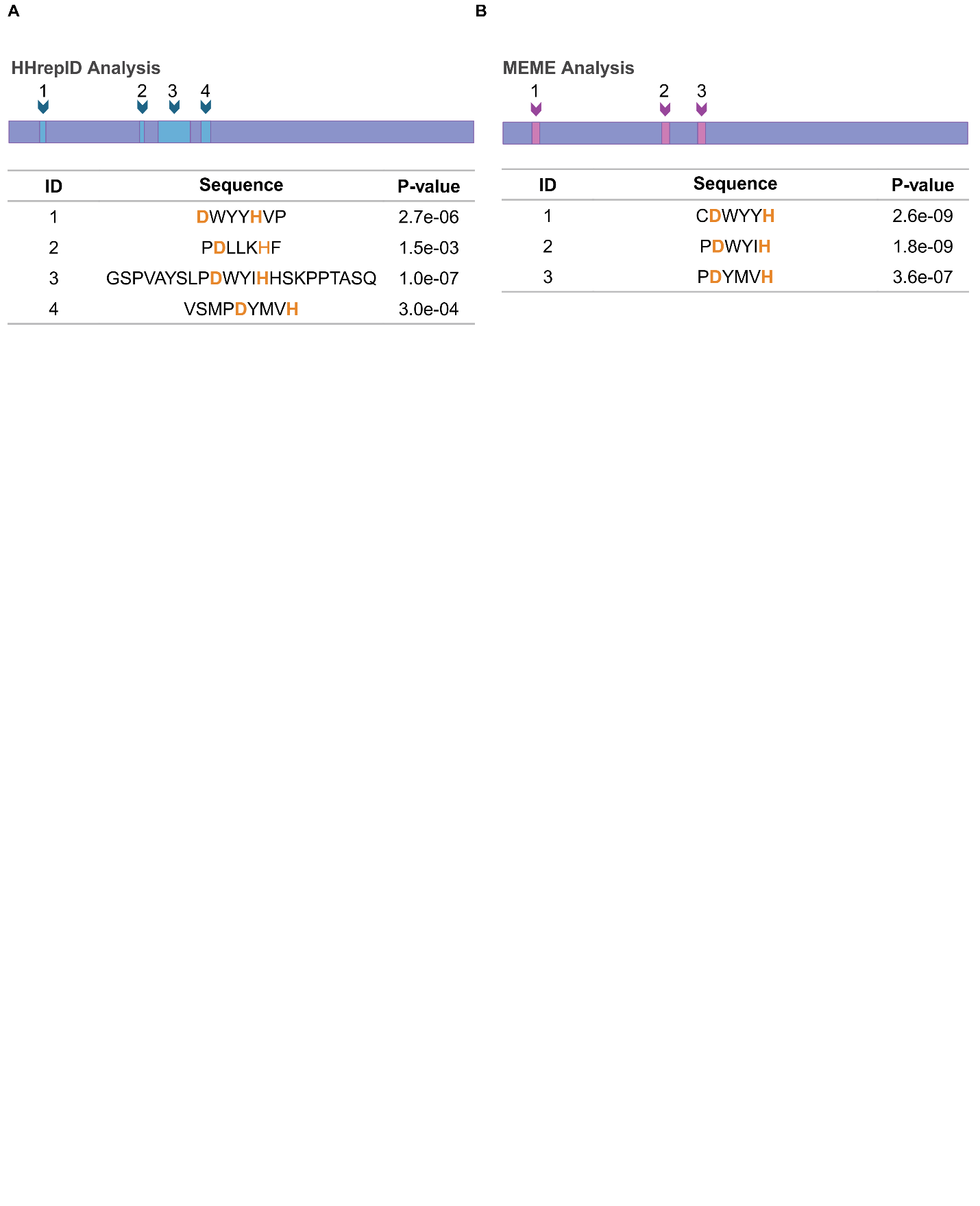
**Figure S5:** **De novo identification of internal repeats in C7orf57 primary sequence**. (A) HHrepID analysis reveals four short internal repeats at the N-terminus of C7orf57 sequence. (B) MEME analysis reveals three repeats that are also found in HHrepID analysis.

**Table S1 Oligonucleotides used in this study**

| **Oligonucleotides** | | |
| --- | --- | --- |
| Human ACTB RT-qPCR forward primer | Integrated DNA technologies | 5’-CATCCACGAAACTACCTTCAACTCC-3' |
| Human ACTB RT-qPCR reverse primer | Integrated DNA technologies | 5’-GAGCCGCCGATCCACACG-3’ |
| Human SEPT9 RT-qPCR forward primer | Integrated DNA technologies | 5’-TCAACATCGTCCCTGTCATCG-3' |
| Human SEPT9 RT-qPCR reverse primer | Integrated DNA technologies | 5’-AAATTCCTTCTGGGGGTACACG-3' |
| Human PSRC1 RT-qPCR forward primer | Integrated DNA technologies | 5’-TGTCAACTCGAAAAGGGCTTCC-3' |
| Human PSRC1 RT-qPCR reverse primer | Integrated DNA technologies | 5’-TGGGAAACAGGAACCTTGTGTC-3' |
| Human BBS5 RT-qPCR forward primer | Integrated DNA technologies | 5’-TGCCAGTCCCATATTTGGAGTTG-3' |
| Human BBS5 RT-qPCR reverse primer | Integrated DNA technologies | 5’-AAAGCATCCGTGTGACCATCAG-3' |
| Human NUBP2 RT-qPCR forward primer | Integrated DNA technologies | 5’-TAGAAGCCCTGCGTCCCTAC-3' |
| Human NUBP2 RT-qPCR reverse primer | Integrated DNA technologies | 5’-TGAAGCCGCTCATATTCTCCAC-3' |
| Human CCDC34 RT-qPCR forward primer | Integrated DNA technologies | 5’-ACGTGGATGAAGATGCCCATG-3' |
| Human CCDC34 RT-qPCR reverse primer | Integrated DNA technologies | 5’-GCGCACCTGTTTCTGTTCTTC-3' |
| Human CKAP2 RT-qPCR forward primer | Integrated DNA technologies | 5’-TGACCAGCGAAGACATACTGC-3' |
| Human CKAP2 RT-qPCR reverse primer | Integrated DNA technologies | 5’-CACTCTTCCTTTGCCAGCTTTC-3' |
| Human DPYSL2 RT-qPCR forward primer | Integrated DNA technologies | 5’-ATCGCTTCCAGCTAACGGATTG-3' |
| Human DPYSL2 RT-qPCR reverse primer | Integrated DNA technologies | 5’-TGCTGCTCCTCTGCAATGATG-3' |
| Human INVS RT-qPCR forward primer | Integrated DNA technologies | 5’-ACAAGGTCAGAAAAGCCTTCCG-3' |
| Human INVS RT-qPCR reverse primer | Integrated DNA technologies | 5’-TCCGCCTTCCTTTTTGCTGTTC-3' |
| Human WDR34 RT-qPCR forward primer | Integrated DNA technologies | 5’-GTTTGATGGCTTCGAGGTGAAC-3' |
| Human WDR34 RT-qPCR reverse primer | Integrated DNA technologies | 5’-AGGAGATGCTGGTCACATGC-3' |
| Human TBC1D31 RT-qPCR forward primer | Integrated DNA technologies | 5’-TGTAACCAGAGATGGCCGAATC-3' |
| Human TBC1D31 RT-qPCR reverse primer | Integrated DNA technologies | 5’-TCGAACTTTAGTGGGCATCTGG-3' |
| Human MPP9 RT-qPCR forward primer | Integrated DNA technologies | 5’-AGCAGAACCAACCAAAGAGAGC-3' |
| Human MPP9 RT-qPCR reverse primer | Integrated DNA technologies | 5’-AAAGCACGAGGATGTGTAGCAG-3' |
| Human USP21 RT-qPCR forward primer | Integrated DNA technologies | 5’-GTCTCTGCGGGATTGTTTCAAC-3' |
| Human USP21 RT-qPCR reverse primer | Integrated DNA technologies | 5’-TTCAGATGGAGCACGAGGATTC-3' |
| Human AKT1 RT-qPCR forward primer | Integrated DNA technologies | 5’-TTGCTTTCAGGGCTGCTCAAG-3' |
| Human AKT1 RT-qPCR reverse primer | Integrated DNA technologies | 5’-ATACCGGCAAAGAAGCGATGC-3' |
| Human FAM161A RT-qPCR forward primer | Integrated DNA technologies | 5’-CGAACGCGAAGACCCCTTAAAG-3' |
| Human FAM161A RT-qPCR reverse primer | Integrated DNA technologies | 5’-TGGTGTTCAAATCAGCCGATGC-3' |
| Human TBCCD1 RT-qPCR forward primer | Integrated DNA technologies | 5’-AACTGCATGTCTGAGGAGGAAG-3' |
| Human TBCCD1 RT-qPCR reverse primer | Integrated DNA technologies | 5’-GCCACTCTTCGCCAATCAAAG-3' |
| Human CKAP2L RT-qPCR forward primer | Integrated DNA technologies | 5’-TGGGGCAACACCAATACAAGAG-3' |
| Human CKAP2L RT-qPCR reverse primer | Integrated DNA technologies | 5’-ATCTTCTTGGCCAGCTCTTCC-3' |
| Human DCDC2 RT-qPCR forward primer | Integrated DNA technologies | 5’-GGCATTTTCAAAGCTGGAGCAG-3' |
| Human DCDC2 RT-qPCR reverse primer | Integrated DNA technologies | 5’-ATTTCTGCTGGCCTCTGATCG-3' |
| Human HYLS1 RT-qPCR forward primer | Integrated DNA technologies | 5’-TTTGTCGAGCAGAACCCCAATC-3' |
| Human HYLS1 RT-qPCR reverse primer | Integrated DNA technologies | 5’-TTGCAAGGTCACAACGAACACC-3' |
| Human SUFU RT-qPCR forward primer | Integrated DNA technologies | 5’-AACATGGCCCGCAGAGTTAATG-3' |
| Human SUFU RT-qPCR reverse primer | Integrated DNA technologies | 5’-ACTGTTATCCAAAGGGCTGTGC-3' |
| siPool #1 targeting human SEPT9 | siTools Biotech | NA |
| siPool #2 targeting human SEPT9 | siTools Biotech | NA |
| siPool #1 targeting human PSRC1 | siTools Biotech | NA |
| siPool #2 targeting human PSRC1 | siTools Biotech | NA |
| siPool #1 targeting human BBS5 | siTools Biotech | NA |
| siPool #1 targeting human NUBP2 | siTools Biotech | NA |
| siPool #1 targeting human CCDC34 | siTools Biotech | NA |
| siPool #1 targeting human CKAP2 | siTools Biotech | NA |
| siPool #1 targeting human DPYSL2 | siTools Biotech | NA |
| siPool #1 targeting human INVS | siTools Biotech | NA |
| siPool #1 targeting human WDR34 | siTools Biotech | NA |
| siPool #1 targeting human TBC1D31 | siTools Biotech | NA |
| siPool #1 targeting human MPP9 | siTools Biotech | NA |
| siPool #1 targeting human USP21 | siTools Biotech | NA |
| siPool #1 targeting human AKT1 | siTools Biotech | NA |
| siPool #1 targeting human FAM161A | siTools Biotech | NA |
| siPool #1 targeting human TBCCD1 | siTools Biotech | NA |
| siPool #1 targeting human CKAP2L | siTools Biotech | NA |
| siPool #1 targeting human DCDC2 | siTools Biotech | NA |
| siPool #1 targeting human HYLS1 | siTools Biotech | NA |
| siPool #1 targeting human SUFU | siTools Biotech | NA |

**Table S2 Recombinant DNAs used in this study**

| Recombinant DNA | | |
| --- | --- | --- |
| Human C7orf57 in pcDNA3.1-C-(k)DYK vector | Genscript | OHu01369 |
| Human CCDC34 in pcDNA3.1-C-(k)DYK vector | Genscript | OHu04498 |
| Human PSRC1 in pcDNA3.1-C-(k)DYK vector | Genscript | OHu16967 |
| Human CRYL1 in pcDNA3.1-C-(k)DYK vector | Genscript | OHu07194 |
| Human GMPPB in pcDNA3.1-C-(k)DYK vector | Genscript | OHu20446 |
| Human WDR5B in pcDNA3.1-C-(k)DYK vector | Genscript | OHu29471 |
| Human CTDSP1 in pcDNA3.1-C-(k)DYK vector | Genscript | OHu32153 |
| Human PHGDH in pcDNA3.1-C-(k)DYK vector | Genscript | OHu19607 |
| Human EFHC2 in pcDNA3.1-C-(k)DYK vector | Genscript | OHu19450 |
| Human PYCR3 in pcDNA3.1-C-(k)DYK vector | Genscript | OHu01383 |
| DCLK1 (iso1) - NanoLuc Fusion vector | Addgene | Plasmid# 237193 |
| pMD2.G | Addgene | Plasmid# 12259 |
| psPAX2 | Addgene | Plasmid# 12260 |
| pcDNA3.1 mycBioID | Addgene | Plasmid# 35700 |
| Flag tagged human C7orf57 in pCDH-CMV-MCS-EF1-Neo vector | This paper | NA |
| Flag tagged human CCDC34 in pCDH-CMV-MCS-EF1-Neo vector | This paper | NA |
| Flag tagged human PSRC1 in pCDH-CMV-MCS-EF1-Neo vector | This paper | NA |
| Flag tagged human CRYL1 in pCDH-CMV-MCS-EF1-Neo vector | This paper | NA |
| Flag tagged human GMPPB in pCDH-CMV-MCS-EF1-Neo vector | This paper | NA |
| Flag tagged human WDR5B in pCDH-CMV-MCS-EF1-Neo vector | This paper | NA |
| Flag tagged human CTDSP1 in pCDH-CMV-MCS-EF1-Neo vector | This paper | NA |
| Flag tagged human PHGDH in pCDH-CMV-MCS-EF1-Neo vector | This paper | NA |
| Flag tagged human DCLK1 in pCDH-CMV-MCS-EF1-Neo vector | This paper | NA |
| Flag tagged human EFHC2 in pCDH-CMV-MCS-EF1-Neo vector | This paper | NA |
| Flag tagged human PYCR3 in pCDH-CMV-MCS-EF1-Neo vector | This paper | NA |
| mCherry tagged human CEP41 in pCDH-CMV-MCS-EF1-Neo vector | This paper | NA |
| MycBioID in pCDH-CMV-MCS-EF1-Neo vector | This paper | NA |
| Human CEP41-mycBioID fusion gene in pCDH-CMV-MCS-EF1-Neo vector | This paper | NA |
